# A GID E3 ligase assembly ubiquitinates an Rsp5 E3 adaptor and regulates plasma membrane transporters

**DOI:** 10.1101/2021.09.02.458684

**Authors:** Christine R. Langlois, Viola Beier, Ozge Karayel, Jakub Chrustowicz, Dawafuti Sherpa, Matthias Mann, Brenda A. Schulman

## Abstract

Cells rapidly remodel their proteomes to align their cellular metabolism to environmental conditions. Ubiquitin E3 ligases enable this response, by facilitating rapid and reversible changes to protein stability, localization, or interaction partners. In *S. cerevisiae*, the GID E3 ligase regulates the switch from gluconeogenic to glycolytic conditions through induction and incorporation of the substrate receptor subunit Gid4, which promotes the degradation of gluconeogenic enzymes. Here, we show an alternative substrate receptor, Gid10, which is induced in response to changes in temperature, osmolarity and nutrient availability, and regulates the ART-Rsp5 pathway. Art2 levels are elevated upon *GID10* deletion, a crystal structure shows the basis for Gid10-Art2 interactions, and Gid10 directs a GID E3 ligase complex to ubiquitinate Art2. We also find that the GID E3 ligase affects the flux of plasma membrane nutrient transporters during heat stress. The data reveal GID as a system of E3 ligases with metabolic regulatory functions outside of glycolysis and gluconeogenesis, controlled by distinct stress-specific substrate receptors.

## Introduction

The ubiquitin system is an integral part of cellular responses to environmental changes. The post-translational modification of proteins with ubiquitin modulates virtually every cellular pathway and all aspects of protein fate, including gene expression, and protein activity, stability, localization and binding partners. Because of their wide-reaching effects, substrate selection by E3 ubiquitin ligases must be strictly controlled to maintain cellular homeostasis upon environmental perturbations. Indeed, the cell simultaneously employs several control mechanisms to ensure faithful selection of E3 ligase substrates, but how these mechanisms are coordinated across cellular pathways remains poorly understood.

The transfer of one or more ubiquitins to substrate proteins requires a hierarchical pathway, in which ubiquitin is first activated by an E1 ubiquitin activating enzyme, and then transferred to an E2 ubiquitin conjugating enzyme, which together with an E3 ubiquitin ligase covalently attaches ubiquitin to the substrate [1]. The two largest E3 ligase families are the HECT (*H*omologous to the *E*6AP *C*arboxyl *T*erminus)-type E3 ligases, which first pass the ubiquitin from the E2 to the E3 before substrate attachment, and the RING (*R*eally *I*nteresting *N*ew *G*ene) E3 ligases, which facilitate the direct transfer of ubiquitin from E2 to a remotely bound substrate [2,3].

E3 ligases can encompass substrate binding and a catalytic RING or HECT domain all within a single subunit, or substrates can be recruited through receptor subunits. The best-studied examples of such multi-subunit E3s are the Cullin RING Ligases (CRLs), which have a modular architecture, consisting of separable E3 ligase core and interchangeable substrate receptor (SR) elements, providing a means for linking one catalytic unit to potentially thousands of substrates [4-8]. In a related vein, although many HECT E3 ligases are thought to be single subunit enzymes, some members of the Nedd4 family employ adaptor proteins for substrate selection [9-13]. Although constellations of WW domains in Nedd4-family E3 ligases directly recognize PYx(Y/F) motifs in some substrates, in some cases, an intervening PYx(Y/F)-containing adaptor bridges the E3 ligase and substrate [14-18]. Nedd4-family adaptors can also modulate the E3’s sub-cellular localization, which can promote ubiquitination of specific substrates while sequestering the ligase away from others [19-21]. In the budding yeast *S. cerevisiae*, Rsp5 is the sole Nedd4 family E3 ubiquitin ligase, and its PYx(Y/F)-containing adaptor proteins, termed ARTs (*<u>A</u>*rrestin-*<u>R</u>*elated *<u>T</u>*rafficking adaptors), are best recognized for regulating endocytosis of plasma membrane nutrient transporters to serve metabolic needs [10,22,23]. The ART family consists of 14 such adaptor proteins, in a complex network involving activation in response to specific environmental stimuli [24-26]. Furthermore, Rsp5 contains an intrinsic ubiquitin binding site and in many cases ubiquitination of ART proteins promotes their activity [10,24,27,28], suggesting that adaptor ubiquitination may serve as an additional layer of regulation.

Nutrient signaling originating at the plasma membrane also simultaneously regulates cellular synthesis of metabolites. For example, in the absence of glucose, glucose transporters are rapidly endocytosed, and the cell additionally initiates transcriptional and translational programs to promote gluconeogenesis. When glucose becomes available again, cells rapidly restore glucose transporters to the plasma membrane, terminate gluconeogenesis, and resume the more energetically favorable glycolysis [29-31]. One regulator of this response in *S. cerevisiae* is the multi-protein GID E3 ligase, named for mutations in subunits being *<u>G</u>*lucose *I*nduced degradation *<u>D</u>*eficient. Upon glucose availability following carbon starvation, the GID E3 ligase targets rate-limiting gluconeogenic enzymes, including fructose 1,6-bisphosphatase (Fbp1) and malate dehydrogenase (Mdh2), for degradation [32-35]. Intriguingly, despite the critical function carried out by the GID complex during glycolytic growth, subunits of the GID E3 ligase are dispensable for viability and there is no characterized phenotype of GID deletions [32]. While the function of the GID E3 ligase during the switch from gluconeogenic to glycolytic conditions is relatively well-characterized, several lines of evidence suggest that the GID E3 ligase is competent to regulate additional substrates and metabolic pathways in response to a variety of stressors.

First, the GID complex forms multiple distinct assemblies *in vivo*, with each assembly promoting the targeting of discrete substrates. For example, incorporation of Gid7 into GID^SR4^ results in the formation of a supramolecular chelate assembly (Chelator-GID), uniquely suited to target the oligomeric structure of Fbp1 [36]. Second, for both the Chelator assembly harboring Gid7, or singular versions without Gid7, the GID ligase is expressed as an anticipatory complex (GID^Ant^) in virtually all growth conditions, allowing it to rapidly respond to a shift in conditions. GID^Ant^ is comprised of the scaffolding subunits Gid1, Gid5, and Gid8, as well as the RING-like domain containing subunits Gid2 and Gid9. Following a shift in environmental conditions, GID^Ant^ is activated by the binding of an SR to form GID^SR^ [34,35,37]. Third, GID^Ant^ can bind multiple substrate receptors: Gid4, Gid10, and the recently identified Gid11 [38,39]. For example, during the switch from gluconeogenic to glycolytic conditions, the substrate receptor Gid4 is induced and binds GID^Ant^, forming the active GID^SR4^, which in turn recruits gluconeogenic enzymes [33,35,37]. In contrast, heat or osmotic shock induces the expression of Gid10 and the formation of the structurally homologous GID^SR10^ [35,39], the targets of which remain unknown.

The molecular mechanisms underlying coordination of various E3 ligase pathways in response to environmental changes remain poorly understood. To explore these questions, we use the GID E3 ligase as a model multi-functional metabolic regulator. We characterize the regulation of expression of the SRs Gid4 and Gid10. Each SR is transiently induced under distinct environmental conditions, turned-over in a manner which depends on itself, and can influence binding of the other SR. Furthermore, using rapid and high-throughput data independent acquisition (DIA) mass spectrometry (MS)-based proteomics analysis [40], we identify the ART-Rsp5 network as a novel regulatory target of GID^SR10^, demonstrating cross-talk between the two E3 ligase pathways.

## Results

### Gid10 has hallmark features of a GID E3 substrate receptor *in vivo*

Previous studies suggested that Gid10 could be a SR of the GID E3 ligase. For example, it has been shown that both Gid4 and Gid10 bind the GID^Ant^ scaffolding subunit Gid5 [35,37,39]. In addition, a high resolution cryo-EM structure of GID^SR4^ showed Gid4 binding a concave surface of Gid5, through key interactions mediated by its C-terminal tail, and a low resolution structure demonstrated that Gid10 forms a homologous complex [35]. Consistent with this, a yeast two-hybrid analysis confirmed that both Gid10 and Gid4 bind directly to Gid5 (Fig 1A). To investigate if the same intermolecular interactions are required for Gid4 and Gid10 binding, we probed the effect of structure-based mutants in the Gid5-SR binding interface. While Gid4 and Gid10 were able to bind WT GID^Ant^ to a similar extent, binding was significantly abrogated to GID^Ant^ containing Gid5 point mutations (Gid5^W606A, Y613A, Q649A^) on the concave binding surface, which also disrupts ubiquitination by GID^SR4^ [35] (Fig 1B). Furthermore, deletion of the C-terminal residues in Gid4 or Gid10 also significantly reduced the binding of each SR to GID^Ant^ (Fig 1B), indicating that Gid4 and Gid10 bind to the same surface on Gid5 through homologous residues on each SR.

**Figure 1.**
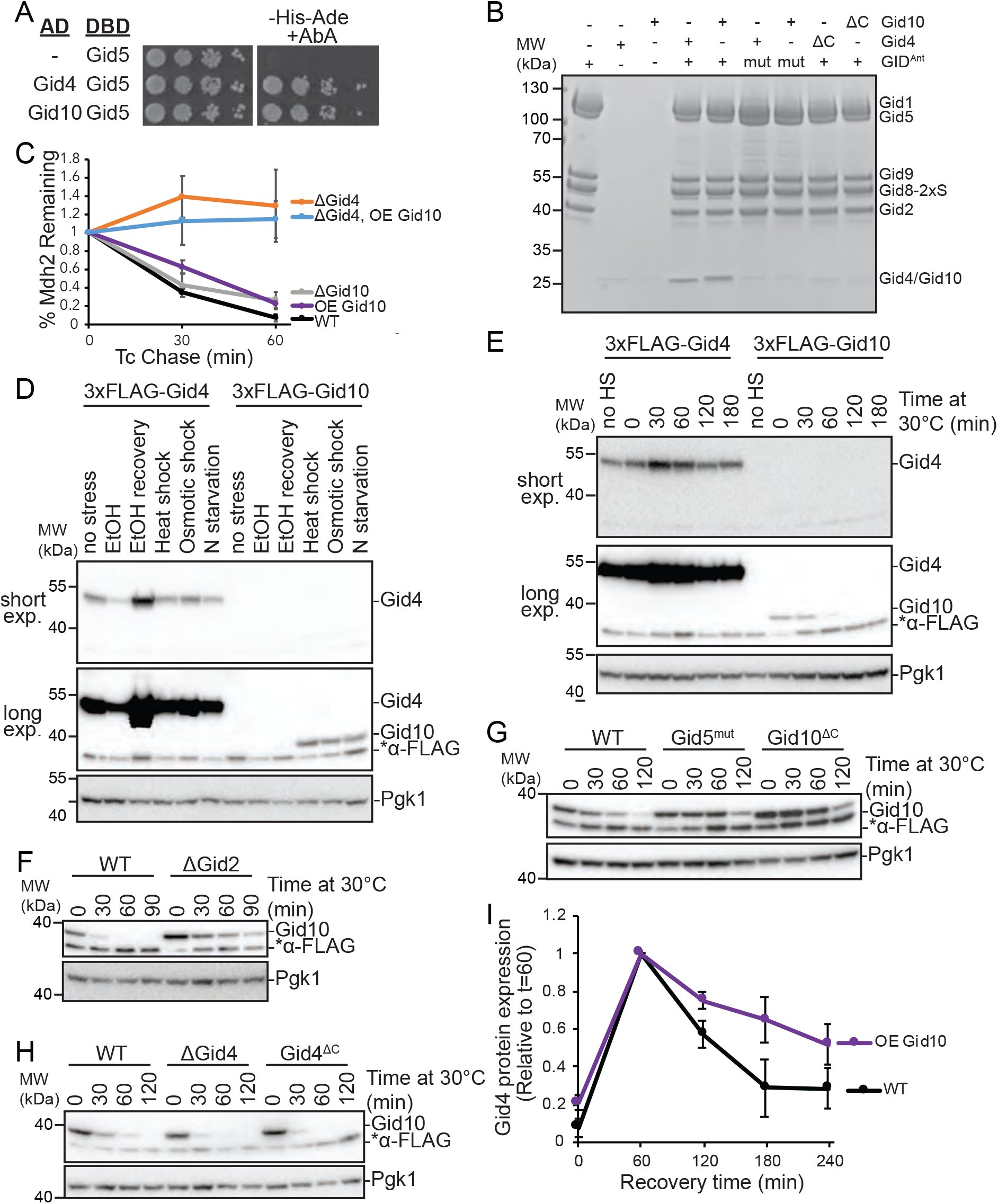
Gid4 and Gid10 expression is regulated by the GID E3 ligase. A) Yeast two-hybrid interactions between SR-Gal4 activation domain (AD) and Gid5-DNA binding domain (DBD). Growth on -His-Ade+Aureobasidin A (AbA) is indicative of an interaction between the two test proteins. Spots represent 1:5 serial dilutions B) Strep-Tactin pull down of GID^Ant^ (strep-tagged at Gid8 C-terminus) probing binding of Gid10^57-292^ and Gid4^117-362^ to the complex visualized with Coomassie-stained SDS-PAGE. The experiment was performed with WT and C-terminal deletion (ΔC) of the substrate receptors (Δ289-292 and Δ359-362 for Gid10 and Gid4, respectively) andWT and mutant (mut, Gid5^W606A/Y613A/Q649A^) GID^Ant^. C) Tetracycline reference-based chase performed during transition from ethanol to glucose media with wildtype, ΔGid4 and ΔGid10 strains, and in wildtype and ΔGid4 strains overexpressing (OE) Gid10. Points represent mean, error bars represent standard deviation (n>3). D) Lysates from yeast strains expressing endogenously tagged 3xFLAG-Gid4 or 3x-FLAG-Gid10 that were grown in SD complete at 30°C (no stress), SE complete for 19 hours (EtOH), SD complete for 1 hour following 19 hour ethanol treatment (EtOH recovery), 42°C for 1 hour (heat shock), SD complete supplemented with 0.5M NaCl for 1 hour (osmotic shock), or SD-N for 1 hour (N-starvation) were run on an SDS-PAGE gel and immunoblotted with αFLAG (two exposures from the same gel are shown) and αPGK. E) Lysates from yeast strains expressing endogenously tagged 3xFLAG-Gid4 or 3xFLAG-Gid10 that were grown at 42°C for one hour, and then returned to 30°C for the indicated timepoints were run on an SDS-PAGE gel and immunoblotted with αFLAG (two exposures from the same gel are shown) and αPGK. F) Lysates from wildtype and ΔGid2 yeast strains expressing endogenously tagged 3xFLAG-Gid10 that were grown at 42°C for one hour, and then returned to 30°C for the indicated timepoints were run on an SDS-PAGE gel and immunoblotted with αFLAG and αPGK. G) Lysates from wildtype, Gid5^W606A,Y613A,Q649A^, and Gid10^ΔC^ strains expressing endogenously tagged 3xFLAG-Gid10 that were grown at 42°C for one hour, and then returned to 30°C for the indicated timepoints were run on an SDS-PAGE gel and immunoblotted with αFLAG and αPGK. H) Lysates from wildtype, ΔGid4, and Gid4^ΔC^ strains expressing endogenously tagged 3xFLAG-Gid10 that were grown at 42°C for one hour, and then returned to 30°C for the indicated timepoints were run on an SDS-PAGE gel and immunoblotted with αFLAG and αPGK. I) Wildtype and Gid10 overexpressing yeast strains expressing endogenously tagged Gid4 were grown for 19 hours in YPE, and transitioned to YPD for the indicated timepoints. Lysates were run on an SDS-PAGE gel and immunoblotted with αFLAG and αPGK. Points represent mean, error bars represent standard deviation (n>3).

Gid4 and Gid10 share many sequence and structural elements and might carry out redundant functions in the cell. Indeed, GID^SR10^ is capable of ubiquitinating Mdh2 *in vitro*, albeit to a lesser extent than GID^SR4^ [35]. However, Gid10 is unable to substitute for loss of Gid4 *in vivo* to promote Mdh2 or Fbp1 degradation, even when placed under the control of the Gid4 promoter [39]. Because ubiquitination of Mdh2 *in vitro* by GID^SR10^ is less efficient than by GID^SR4^, higher levels of Gid10 might be required to compensate for lack of Gid4 *in vivo*. To test this, we employed the promoter reference technique, which allows the fate of existing proteins to be monitored without the use of global transcription or translation inhibitors [33,41]. During the switch between gluconeogenic and glycolytic conditions, the GID^SR4^ substrates Mdh2 and Fbp1 are significantly stabilized in a Gid4 deletion strain, but not in a Gid10 deletion strain (Fig 1C, EV1A), in agreement with previously published results [34,39]. In addition, constitutive overexpression of Gid10 from the Tdh3 promoter alone did not significantly alter the rate of Mdh2 or Fbp1 degradation, and could not compensate for loss of Gid4 (Fig 1C, EV1B). Thus, even overexpressed Gid10 is not competent to promote recognition or degradation of Gid4 substrates during carbon recovery *in vivo*.

While Gid10 protein levels are not induced during carbon starvation or recovery, they are transiently induced in response to a variety of other stress conditions, including heat shock, osmotic shock, and amino acid and nitrogen starvation (Fig 1D, EV1C-H) [35,39,42]. Interestingly, while Gid10 is induced during heat shock, Gid4 is transiently induced during recovery from heat shock (Fig 1E), suggesting complementary roles of the two SRs during stress and recovery. To gain a better understanding of how the transient expression of SRs is regulated, we first examined the requirements of the GID complex for SR turnover. All of the subunits of GID^Ant^ were previously shown to be required for Gid4 turnover [43], and thus we hypothesized that SR degradation may be triggered after binding GID^Ant^. Indeed, both Gid4 and Gid10 are stabilized when the RING-like containing subunit Gid2 is deleted (Fig 1F, EV1I). Furthermore, Gid4 and Gid10 are also stabilized when their ability to bind to GID^Ant^ via Gid5 is impaired (Fig 1G, EV1J), demonstrating that both GID complex activity and SR binding are required for SR turnover.

Gid10 protein expression is significantly lower than Gid4 under all tested conditions (Fig 1D-E). Therefore, if the two SRs compete for binding to GID^Ant^ via Gid5, absence of Gid4 should significantly affect the kinetics of Gid10 turnover, but absence of Gid10 should have little to no effect on Gid4 turnover. Indeed, Gid10 turnover during heat shock recovery is accelerated by either a deletion of Gid4 or expression of a GID^Ant^-binding impaired Gid4 (Fig 1H). In contrast, Gid4 turnover during carbon recovery is largely unaffected by the absence of Gid10 (Fig EV1I); when Gid10 is constitutively over-expressed, and therefore better able to compete for GID^Ant^ binding, Gid4 turnover during recovery from carbon starvation is delayed (Fig 1I). Taken together, these data are consistent with a model in which the SRs can compete with each other for access to GID^Ant^ and that binding is a prerequisite for SR turnover, suggesting that the SRs may be auto-ubiquitinated when bound to GID^Ant^.

### Art2 is a regulatory target of GID^SR10^

Biological functions for the Gid10 protein remain elusive. To identify its regulatory targets, we employed a systems-wide single-run DIA-based proteomics approach [40] during heat stress, when Gid10 is maximally expressed. Following a one-hour heat shock at 42°C, only two proteins were significantly upregulated in a Gid10 deletion strain, compared to wild type: Art2, a member of the α-arrestin family, and Nhp10, a member of the INO80 chromatin remodeling complex (p value<0.01 and fold change>4, Fig 2A) [44,45]. Importantly, regulation of both proteins was Gid10-specific as their abundance did not change in a Gid4 deletion strain (Fig 2B). To determine if Art2 or Nhp10 might also be regulated under other growth conditions where the GID E3 ligase is known to be active, we reanalyzed our previously published data set characterizing GID-dependent protein regulation during recovery from ethanol starvation [40]. We selected for proteins that contain a proline in position 2 or 3, and are significantly upregulated during growth in ethanol, compared to glucose, and downregulated during recovery compared to ethanol. Interestingly, Art2 also appears to be regulated by Gid2 (although this did not reach statistical significance), but not by Gid4, during recovery from ethanol starvation (Fig EV2).

**Figure 2.**
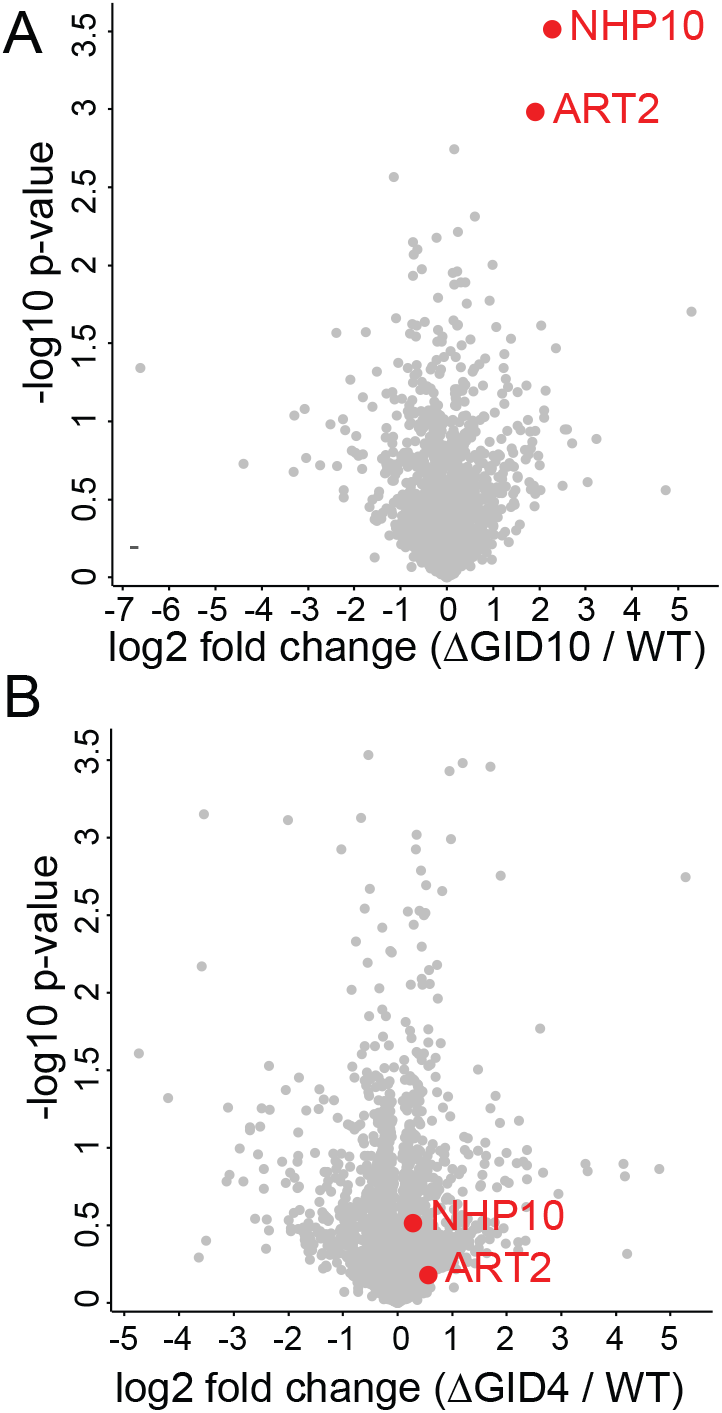
Art2 is upregulated in the absence of Gid10 during heat shock. A) Volcano plot of the (−log10) p values vs. the log2 protein abundance differences between Gid10 null yeast vs. WT. Red dots indicate significantly different proteins, determined based on p value < 0.01 and at least fourfold change. B) Volcano plot of the (−log10) p values vs. the log2 protein abundance differences between Gid4 null yeast vs. WT. Red dots indicate significantly different proteins in the comparison of Gid10 null yeast vs. WT shown in A.

GID^SR4^ is an N-degron E3 ligase that recognizes substrates with an N-terminal proline, although peptides with other N-terminal residues have been shown to bind Gid4 or Gid10 with lower affinity [32,33,39,46,47]. On this basis, we probed the potential for Gid10 interaction to bind the Art2 or Nhp10 N-terminal sequences by yeast two-hybrid. Gid10 was efficiently bound the Art2, but not the Nhp10, N-terminus (Fig 3A, EV3A). Moreover, the Gid10-Art2 interaction is Gid10-specific as we did not observe an interaction between Gid4 and the Art2 N-terminus (Fig 3A), and dependent on the N-terminal proline of Art2 (Fig 3B). In contrast, Gid4, but not Gid10, interacted with the N-terminus of the classic Gid4 substrate Mdh2 (Fig 3A), suggesting that Gid10 and Gid4 indeed prefer discrete regulatory targets.

**Figure 3.**
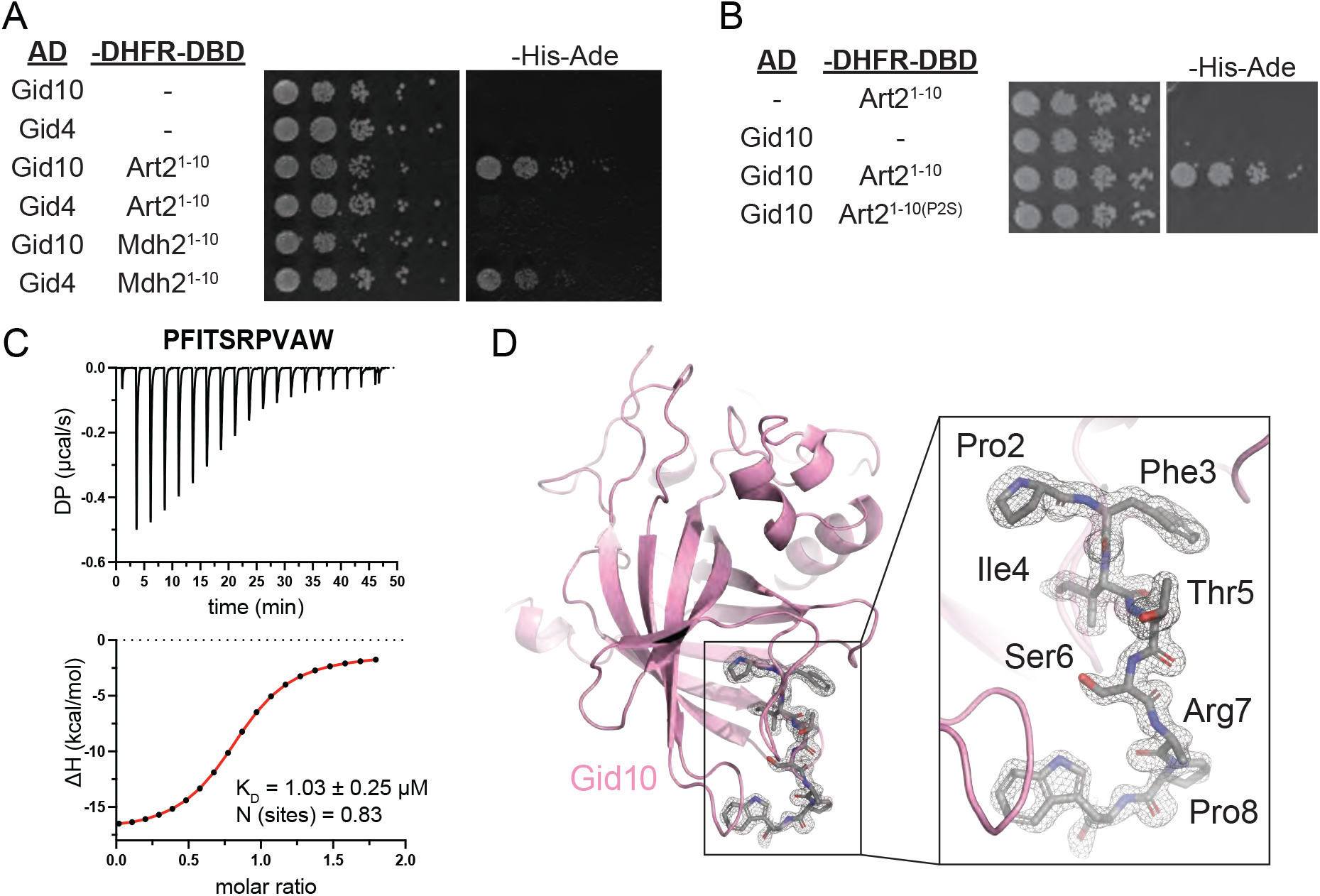
Art2 binds to Gid10 via its N-terminal proline. A) Yeast two-hybrid between SR-Gal4 activation domain (AD) and substrate degrons fused to DHFR-DNA binding domain (-DHFR-DBD). Growth on -His-Ade is indicative of an interaction between the two test proteins. Spots represent 1:5 serial dilutions. B) Yeast two-hybrid between Gid10-Gal4-AD and the Art2^WT^ or Art2^P2S^ degron fused to DHFR-DBD. Growth on -His-Ade is indicative of an interaction between the two test proteins. Spots represent 1:5 serial dilutions. C) Isothermal titration calorimetry (ITC) binding assay to quantify affinity (KD) of Art2^2-9^ degron for Gid10^57-292^ substrate-binding domain. The raw ITC results (top) were integrated to calculate the amount of heat released (ΔH) during every injection of a peptide and plotted as a function of peptide:protein molar ratio (bottom). Fitting of the obtained data points to the binding model served to determine KD and stoichiometry of the binding reaction (N). D) 1.3 Å-resolution crystal structure of Gid10^65-284^ substrate-binding domain (pink cartoon) in complex with Art2^2-8^ degron (grey sticks, C-terminal Trp was attached to accurately measure peptide concentration). The grey mesh represents electron density corresponding to the Art2 peptide.

To further confirm Gid10 binding to the Art2 N-terminus, we quantified the interaction using isothermal titration calorimetry. The putative Gid10 substrate binding domain (Gid10^Δ1-56^) bound the first 9 amino acids of Art2 (PFITSRPVA) synthesized with a C-terminal tryptophan to enable concentration determination based on extinction coefficient at 280 nm, with a KD of 1.03 μM (Fig 3C). Notably, this is two-fold higher affinity than any published peptide examined for interaction with Gid4 [46,47].

To determine the molecular mechanism of this interaction, we determined a crystal structure showing degron recognition by Gid10. A co-crystal structure of Gid10^Δ1-56^ bound to a peptide corresponding to the first seven residues of Art2 (PFITSRP, plus a tryptophan at the C-terminus) showed that Gid10 resembles Gid4 in forming a β-barrel with several helical insertions (Fig 3D). In addition, the binding pocket residues on Gid4 and Gid10, the trajectory of the Fbp1 or Art2 degron in the Gid4 or Gid10 binding pocket, respectively, and the position of the N-terminal proline were strikingly similar (Fig EV3B-C). The structures of the two SRs are nearly identical, with an RMSD of 0.87 Å (Fig EV3B). However, Gid10’s interaction with the Art2 sequence is far more extensive than Gid4 interactions with the Fbp1 degron in the context of Chelator-GID^SR4^, or human Gid4 bound to Pro/N-degron peptides [36,46]. All seven residues of the Art2 peptide interact with Gid10, explaining the relatively high affinity of this interaction.

To further characterize Art2 as a GID^SR10^ substrate, we asked if Gid^SR10^ was capable of ubiquitinating Art2. Towards this end, we performed ubiquitination assays using Art2-3xFLAG immunocaptured from yeast lysates, and recombinantly expressed GID^Ant^, GID^SR10^, or GID^SR4^. In this system, GID^SR10^, but not GID^SR4^ or GID^Ant^, was able to efficiently poly-ubiquitinate Art2. In contrast, only GID^SR4^ was capable of ubiquitinating Mdh2 (Fig 4A, B). Moreover, GID^SR10^-mediated ubiquitination of Art2 is dependent on Gid10 binding to Gid^Ant^, as well as the ability of Art2 to bind Gid10 via its N-terminal proline (Fig 4C). To further confirm the ubiquitination activity of GID^SR10^ towards Art2, we used a peptide substrate consisting of residues 2-28 of Art2, which includes an endogenously ubiquitinated lysine at position 26 [48]. In the presence of GID^SR10^, we observed poly-ubiquitination of this peptide substrate, which was dependent on the N-terminal proline. GID^SR4^ also mediated low-level ubiquitination of the peptide substrate (Fig 4D), which suggests the potential to interact in the context of a fully-assembled E3. Taken together, our results indicate that Art2 is a substrate of GID^SR10^.

**Figure 4.**
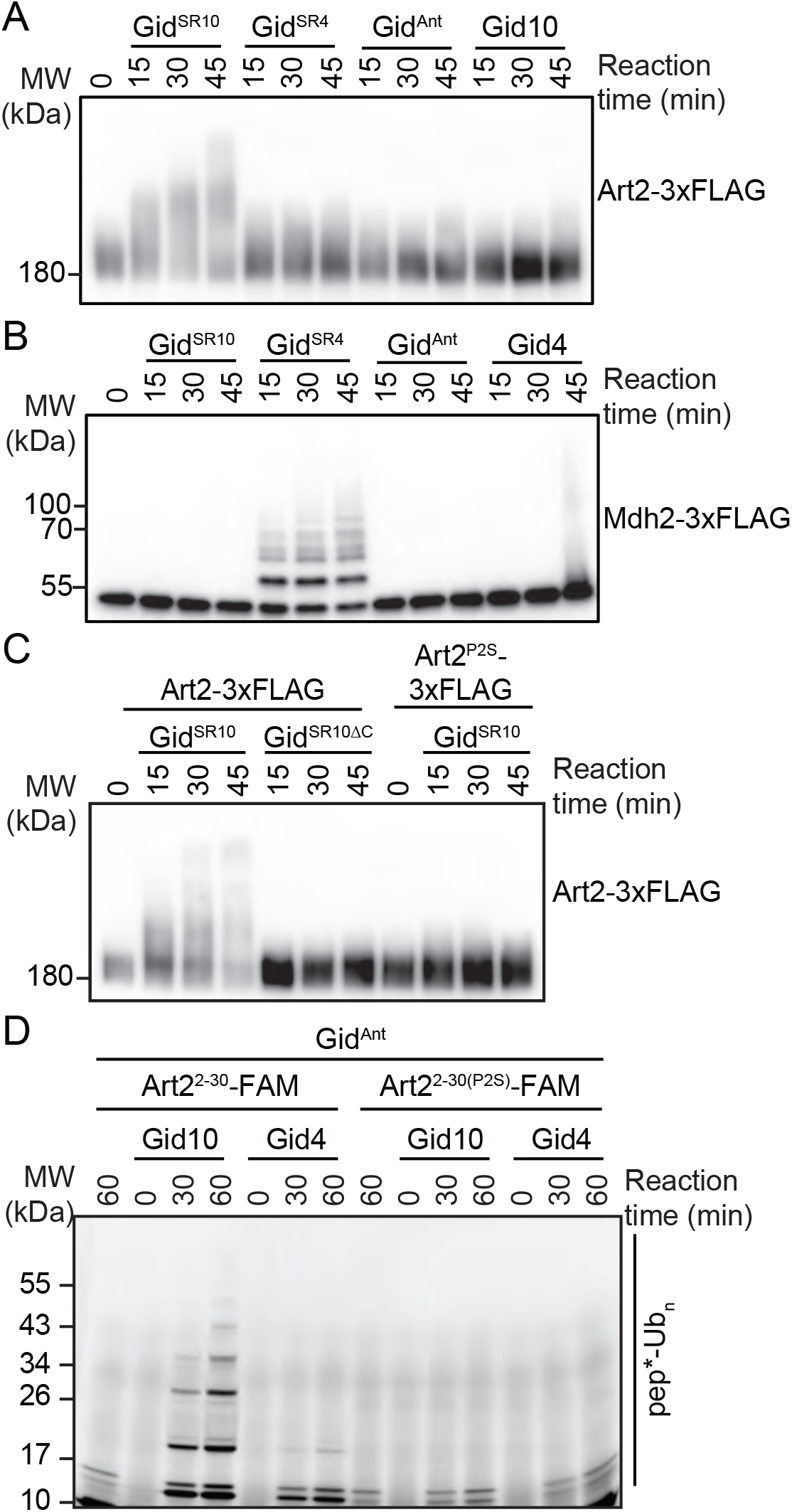
Gid^SR10^ ubiquitinates Art2. A) Art2-3xFLAG was immunocaptured from yeast cells grown in YPD and incubated with Gid^SR10^, Gid^SR4^, Gid^Ant^, or Gid10 for the indicated timepoints. Progress of the reaction was followed by αFLAG immunoblot. B) Mdh2-3xFLAG was immunocaptured from yeast cells following growth in YPE for 19h and incubated with Gid^SR10^, Gid^SR4^, Gid^Ant^, or Gid4 for the indicated timepoints. Progress of the reaction was followed by αFLAG immunoblot. C) Art2-3xFLAG or Art2^P2S^-3xFLAG was in YPD and incubated with Gid^SR10^, or Gid^SR10ΔC^ for the indicated timepoints. Progress of the reaction was followed by αFLAG immunoblot. *D) In vitro* ubiquitination assay probing the ability of Gid10 and Gid4 to promote ubiquitination of Art2^2-28^ N-terminus and its P2S mutant. Progress of the reaction was monitored by fluorescent scan of the gel visualizing the Art2 peptide with FAM appended to its C-terminus (pep*).

### The GID E3 ligase affects flux of plasma membrane nutrient transporters

Art2 is one of a suite of ART adaptors that guides the Nedd4 family E3 ubiquitin ligase Rsp5 to plasma membrane transporters, and directs their selective endocytosis [22,49]. Intriguingly, Rsp5 as well as the α-arrestin Art3 also contain N-terminal prolines. Given that their other substrates contain N-terminal prolines, we tested if Gid10 and/or Gid4 bind the N-terminal sequences of Rsp5 or members of the α-arrestin family. However, our yeast two-hybrid assay revealed only the Gid10-Art2 interaction, highlighting its specificity (Fig EV4A-C).

Compared to other ARTs, the functions of Art2 are relatively poorly characterized, in part because of redundancy with other more functionally dominant family members, and because its relatively large size and protein properties make biochemical analyses challenging.

Nonetheless, Art2 has been associated with endocytosis and vacuolar degradation of the lysine permease, Lyp1, during some environmental perturbations, including amino acid starvation, nitrogen starvation, and cycloheximide treatment [10,26]. Thus, we tested if the GID complex plays a role in Lyp1 import and degradation by examining phenotypes on the toxic lysine analog, thialysine (S-Aminoethyl-l-cysteine). Importantly, both Art2 deletion and an Art2^P2S^ mutant also showed delayed growth on thialysine (Fig 5A), suggesting that the Art2 deletion effect can at least partly be attributed to regulation by the GID E3 ligase. Furthermore, individual deletions of all GID core subunits, with the exception of Gid7, which is dispensable for some substrates [36,38,50], resulted in cellular toxicity during growth on thialysine, even in the absence of an additional stress condition (Fig 5B). The similar phenotypes for all core subunits suggest a role in regulation of Lyp1 receptor localization or activity. Although deletion of *GID10* or *GID4* individually did not cause a noticeable growth defect on thialysine, the double deletion of both substrate receptor subunits, resulted in a defect similar to that observed upon deletion of core GID subunits (Fig 5B) suggesting that there may be some overlap in SR function *in vivo*, consistent with the low level of GID^SR4^ activity seen in the *in vitro* ubiquitination assay (Fig 4D).

**Figure 5.**
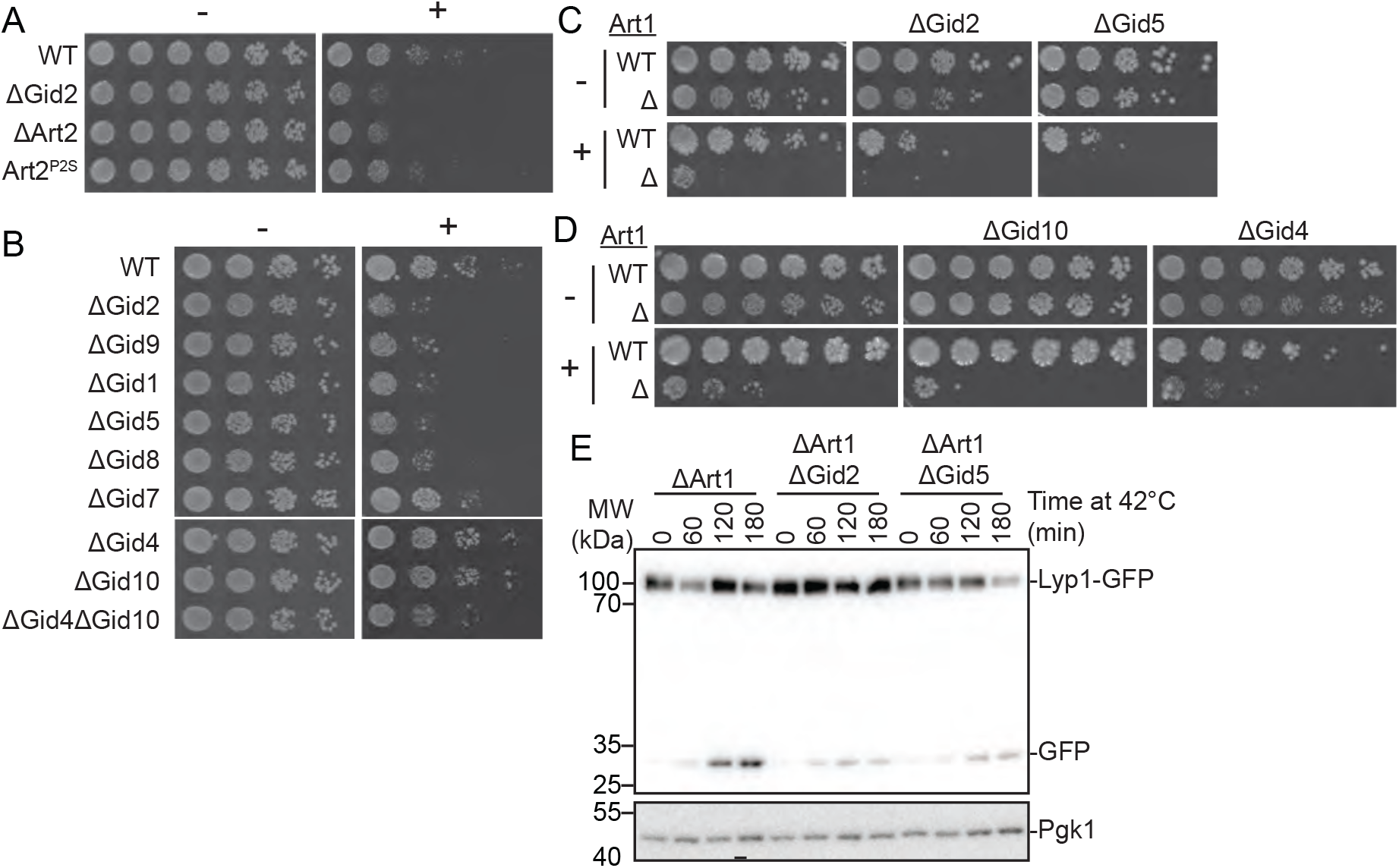
The GID E3 ligase affects flux of plasma membrane nutrient transporters. A) Growth assay of wildtype, ΔGid2, ΔArt2, and Art2^P2S^ yeast strains on SD-Lys (-) and SD-Lys containing 1.0 μg/mL thialysine (+). Spots represent 1:2.6 serial dilutions. B) Growth assay of wildtype yeast or yeast strains containing the indicated deletions on SD-Lys (-) and SD-Lys containing 1.5 μg/mL thialysine (+). Spots represent 1:5 serial dilutions. C) Growth assay of wildtype and ΔArt1 strains containing *GID2* or *GID5* deletions on SD-Lys (-) and SD-Lys containing 1.0 μg/mL thialysine (+). Spots represent 1:5 serial dilutions. D) Growth assay of wildtype or *GID4* deletions on SD-Lys (-) and SD-Lys containing 1.0 μg/mL thialysine (+). Spots represent 1:2.6 serial dilutions. E) ΔArt1 strains containing *GID2* or *GID5* deletions and expressing endogenously tagged Lyp1-GFP were grown at 42°C for the indicated timepoints. Lysates were immunoblotted for αGFP and αPGK.

We used a GFP protection assay to determine if the GID E3 ligase impacts Lyp1 import and degradation. Because GFP is resistant to vacuolar degradation, but proteins fused to it typically are not, the appearance of free GFP on a western blot upon expression of GFP-tagged plasma membrane proteins reflects delivery of the GFP-tagged protein to the vacuole. Performing this assay with Lyp1-GFP showed that Lyp1 import and degradation during heat shock remained unchanged in the GID mutant strains (Fig EV5A). This was surprising, given the effect of these mutations on yeast growth on thialysine. In addition to Art2 regulation of Lyp1, another ART protein, Art1, has also been shown to regulate Lyp1 import and degradation in response to lysine excess, thialysine treatment, and heat stress [10,23,51], suggesting that GID-dependent regulation of Art2 may not be the main mechanism to promote Lyp1 import during heat stress. Indeed, we observe that Lyp1 import and degradation is significantly reduced in an Art1 mutant, compared to wildtype, and a double deletion of Art1 and Art2 is further impaired (Fig EV5B). In addition, an Art1 deletion showed a strong growth defect on thialysine, which was further aggravated by deletion of Art2, while over expression of Art2 completely rescued the growth defect of an Art1 deletion under these conditions (Fig EV5C). Taken together, these data suggest that Art1 is the main regulator of Lyp1 during heat stress, but Art2 also contributes to this regulation.

To further probe this combinatorial regulation, we next investigated the contribution of GID subunits to Lyp1 import in strains containing an *ART1* deletion. In the absence of Art1, deletion of a core subunit resulted in increased toxicity during growth on thialysine, similar to that observed in an *ART1ART2* double deletion (Fig EV5C). Furthermore, deletion of Art1 also resulted in increased toxicity in the Gid10 deletion strain, but not in the Gid4 deletion (Fig EV5D). Moreover, deletion of the GID core subunits Gid2 or Gid5 results in impaired Lyp1 import and degradation in the ΔArt1 background (Fig 5E). Deletion of GID subunits in the context of an *ART1* deletion also resulted in similar defects in degradation of the arginine receptor, Can1, during heat shock (Fig EV5D) demonstrating that the effects of the GID complex are not specific to the Lyp1 receptor, but that the GID E3 ligase facilitates a more general response. Taken together, these data indicate that GID^SR10^, and to a lesser extent GID^SR4^, modulate Art2 function to affect the flux of plasma membrane nutrient transporters.

## Discussion

Here, we demonstrate that the GID E3 ligase is a multifunctional metabolic regulator that incorporates different SRs in response to distinct stresses. We show that Gid10 is a bona fide substrate receptor, by identifying Art2 as a protein that binds Gid10 through specific contacts directed by its N-terminal proline, and is ubiquitinated by GID^SR10^. Furthermore, we identify for the first time a physiological phenotype for the yeast GID complex: increased sensitivity to thialysine, which is dependent on core GID subunits and both SRs.

Through ubiquitination of Art2, the GID E3 could affect the activity of another E3, Rsp5. Indeed, Art2 is a modulator of Rsp5, and we found that GID influences the import and degradation of two Rsp5 targets, the Lyp1 and Can1 receptors, although the mechanism remains elusive. In addition, reversing the activity of Gid2 has been shown to disrupt growth on low-tryptophan media [52], implicating GID in the regulation of even more plasma membrane receptors. Both plasma membrane protein trafficking and Rsp5 interactions are intricately regulated by ubiquitin. Thus, Art2 ubiquitination could impact the ART-Rsp5 network in several ways. First, ubiquitination of Art2 by GID^SR10^ may lead to its deactivation, or promote its degradation. Indeed, the levels of Art2 are modestly increased in a Gid10 deletion strain, which led to its discovery as a substrate. Notably, all previously identified GID E3 ligase substrates have been shown to undergo proteasomal degradation following ubiquitination. Although the deactivation of Art2 would impact its role as an Rsp5 adaptor, the inherent complexity, redundancy and interconnectedness of the ART-Rsp5 network raises the possibility that loss of Art2 might not exclusively result in loss of Rsp5 ubiquitination. Rather, relieved of interaction with Art2, Rsp5 could become more available for interactions with other arrestins, which could compensate for loss of Art2, or alternatively shift the preference to non-Art2-dependent targets. Second, Art2 ubiquitination could modulate its protein-protein interactions or activities. Ubiquitinated Art2 may have a different sub-cellular localization, or affinity to Rsp5, as compared to unmodified Art2. For example, ubiquitin linked to Art2 could engage Rsp5’s ubiquitin binding-exosite [27,28,53,54] to enhance the Art2-Rsp5 interaction, which in turn could shift Rsp5 towards Art2-dependent targets. Moreover, ubiquitination could impact multiple Art2 functions simultaneously. Ubiquitination of Art2 may also selectively affect its ability to interact with its plasma membrane targets, leading to a higher affinity for some targets, but a lower affinity for others. Future studies will be required to identify precisely how post-translational modifications modulate the ART-Rsp5 network, specific mechanisms impacted by GID E3-dependent ubiquitination of Art2, and the molecular details underlying the relationship between the GID E3 and plasma membrane nutrient transporters.

Interestingly, previous studies have also linked the GID E3 ligase to regulation of plasma membrane proteins. First, some GID complex units have been shown to play a role in the Vacuolar Import and Degradation (VID) pathway, which brings proteins to the vacuole for degradation following their endocytosis from the plasma membrane [55,56]. Second, it has been shown that the signals which lead to the degradation of Fbp1 and the hexose transporter Gal2 likely originate from the same biochemical pathway [57]. Third, Gid11 was recently identified as an additional SR of the GID complex. The Gid11 protein also regulates metabolic enzymes involved in amino acid and nucleotide biosynthesis [38], and deletion of *GID11* leads to defects in plasma membrane electron transport [58], suggesting an additional role for Gid11 in regulation of plasma membrane proteins. Importantly, expression of each SR is only induced during a distinct subset of environmental perturbations. Because each environmental change leads to vast, but distinct, remodeling of cellular metabolism, we propose that the GID E3 ligase may have evolved as a common node to regulate nutrient import across the plasma membrane and subsequent cellular synthesis of the necessary metabolites.

Although the GID E3 ligase has long been characterized as functioning during glucose-induced glycolysis, we show that the GID E3 ligase additionally regulates amino acid transporters, and also that there is a GID phenotype linked to amino acid metabolism, similar to effects observed when deleting ART proteins. What advantages might arise from a singular core E3 complex with distinct inputs from and outputs to multiple metabolic pathways? Because environmental changes are often abrupt, the activation of the complex through incorporation of a single protein allows yeast cells to respond rapidly. In addition, the transient expression of SRs, which we show depends on both GID complex activity and SR-GID^Ant^ binding, ensures that substrate selection by GID^SR^ is limited to a pulse following a switch in environmental conditions. Thus, the GID E3 ligase employs rapid “on” and “off” switches which allow it to rapidly and specifically respond to environmental perturbations.

We speculate that the GID E3 may be poised like other post-translational modifying enzymes that serve as metabolic nodes. For example, abundance or paucity of particular metabolites regulate kinase activities of mTOR, which like the GID E3 assembles into different complexes to regulate specific sets of biosynthetic and catabolic processes. Although the GID E3 is not essential in yeast under normal growth conditions, our work suggests there could be distinct requirements under particular environments. Moreover, the GID complex in higher eukaryotes, (termed CTLH - for C-Terminal to LisH) regulates important physiology and is essential for viability [59,60]. Intriguingly, the CTLH complex serves as a regulator of autophagic flux and mTOR signaling, key pathways that integrate cellular responses to environmental changes [61]. While more studies are needed to characterize additional GID regulatory targets, it is clear that the GID E3 ligase is implicated in diverse cellular pathways throughout eukaryotes and serves to enable rapid and robust cellular responses to environmental perturbations.

## Materials and Methods

### Plasmid list

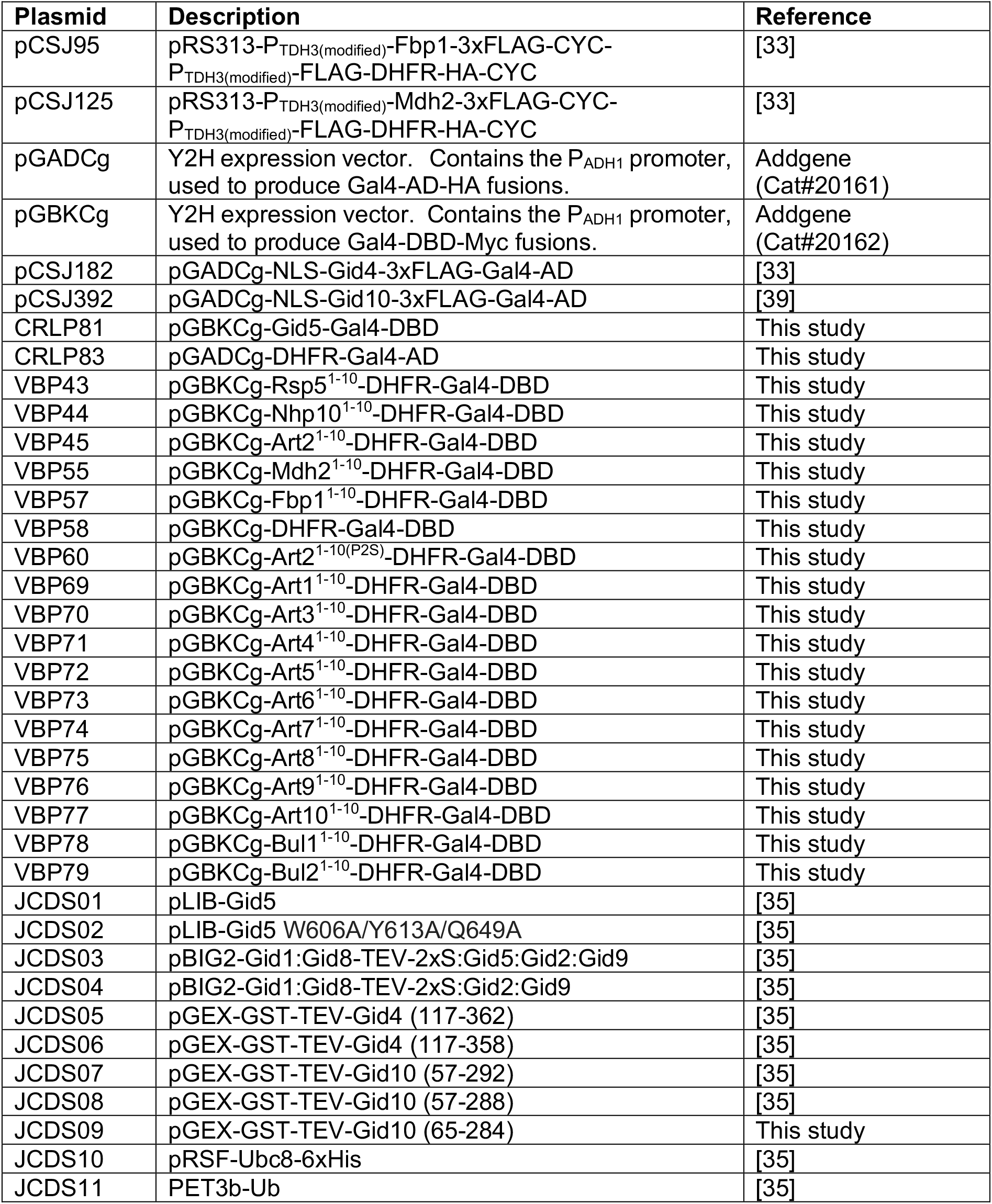

| Plasmid | Description | Reference |
| --- | --- | --- |
| pCSJ95 | pRS313-P<sub>TDH3(modified)</sub>-Fbp1-3xFLAG-CYC-P<sub>TDH3(modified)</sub>-FLAG-DHFR-HA-CYC | [33] |
| pCSJ125 | pRS313-P<sub>TDH3(modified)</sub>-Mdh2-3xFLAG-CYC-P<sub>TDH3(modified)</sub>-FLAG-DHFR-HA-CYC | [33] |
| pGADCg | Y2H expression vector. Contains the P<sub>ADH1</sub> promoter, used to produce Gal4-AD-HA fusions. | Addgene (Cat#20161) |
| pGBKCg | Y2H expression vector. Contains the P<sub>ADH1</sub> promoter, used to produce Gal4-DBD-Myc fusions. | Addgene (Cat#20162) |
| pCSJ182 | pGADCg-NLS-Gid4-3xFLAG-Gal4-AD | [33] |
| pCSJ392 | pGADCg-NLS-Gid10-3xFLAG-Gal4-AD | [39] |
| CRLP81 | pGBKCg-Gid5-Gal4-DBD | This study |
| CRLP83 | pGADCg-DHFR-Gal4-AD | This study |
| VBP43 | pGBKCg-Rsp5<sup>1-10</sup>-DHFR-Gal4-DBD | This study |
| VBP44 | pGBKCg-Nhp10<sup>1-10</sup>-DHFR-Gal4-DBD | This study |
| VBP45 | pGBKCg-Art2<sup>1-10</sup>-DHFR-Gal4-DBD | This study |
| VBP55 | pGBKCg-Mdh2<sup>1-10</sup>-DHFR-Gal4-DBD | This study |
| VBP57 | pGBKCg-Fbp1<sup>1-10</sup>-DHFR-Gal4-DBD | This study |
| VBP58 | pGBKCg-DHFR-Gal4-DBD | This study |
| VBP60 | pGBKCg-Art2<sup>1-10(P2S)</sup>-DHFR-Gal4-DBD | This study |
| VBP69 | pGBKCg-Art1<sup>1-10</sup>-DHFR-Gal4-DBD | This study |
| VBP70 | pGBKCg-Art3<sup>1-10</sup>-DHFR-Gal4-DBD | This study |
| VBP71 | pGBKCg-Art4<sup>1-10</sup>-DHFR-Gal4-DBD | This study |
| VBP72 | pGBKCg-Art5<sup>1-10</sup>-DHFR-Gal4-DBD | This study |
| VBP73 | pGBKCg-Art6<sup>1-10</sup>-DHFR-Gal4-DBD | This study |
| VBP74 | pGBKCg-Art7<sup>1-10</sup>-DHFR-Gal4-DBD | This study |
| VBP75 | pGBKCg-Art8<sup>1-10</sup>-DHFR-Gal4-DBD | This study |
| VBP76 | pGBKCg-Art9<sup>1-10</sup>-DHFR-Gal4-DBD | This study |
| VBP77 | pGBKCg-Art10<sup>1-10</sup>-DHFR-Gal4-DBD | This study |
| VBP78 | pGBKCg-Bul1<sup>1-10</sup>-DHFR-Gal4-DBD | This study |
| VBP79 | pGBKCg-Bul2<sup>1-10</sup>-DHFR-Gal4-DBD | This study |
| JCDS01 | pLIB-Gid5 | [35] |
| JCDS02 | pLIB-Gid5 W606A/Y613A/Q649A | [35] |
| JCDS03 | pBIG2-Gid1:Gid8-TEV-2xS:Gid5:Gid2:Gid9 | [35] |
| JCDS04 | pBIG2-Gid1:Gid8-TEV-2xS:Gid2:Gid9 | [35] |
| JCDS05 | pGEX-GST-TEV-Gid4 (117-362) | [35] |
| JCDS06 | pGEX-GST-TEV-Gid4 (117-358) | [35] |
| JCDS07 | pGEX-GST-TEV-Gid10 (57-292) | [35] |
| JCDS08 | pGEX-GST-TEV-Gid10 (57-288) | [35] |
| JCDS09 | pGEX-GST-TEV-Gid10 (65-284) | This study |
| JCDS10 | pRSF-Ubc8-6xHis | [35] |
| JCDS11 | PET3b-Ub | [35] |

### Yeast strain list

| Strain | Genotype | Reference |
| --- | --- | --- |
| BY4741 | *MATa his3Δ1 leu2Δ0 met15Δ0 ura3Δ0* | Euroscarf (Cat#Y00000) |
| CRLY12 | *BY4741; gid4::KANMX* | [36] |
| CRLY13 | BY4741; *gid5::KANMX* | This study |
| CRLY14 | BY4741; *gid7::KANMX* | [36] |
| CRLY15 | BY4741; *gid8::KANMX* | This study |
| CRLY16 | BY4741; *gid9::KANMX* | This study |
| CRLY17 | BY4741; *gid10::KANMX* | This study |
| CRLY30 | BY4741; *gid2::KANMX* | This study |
| CRLY68 | BY4741; *gid4::3xFLAG-GID4* | [35] |
| CRLY74 | BY4741; *gid10::3xFLAG-GID10* | [35] |
| CRLY186 | BY4741; *gid1::KANMX* | This study |
| CRLY296 | BY4741; *gid10::NATNT2-P<sub>GPD</sub>-GID10* | This study |
| CRLY298 | BY4741; *gid4::KANMX gid10::NATNT2-pGPD-GID10* | This study |
| CRLY301 | BY4741; *gid4::KANMX gid10::3xFLAG-GID10* | This study |
| CRLY314 | BY4741; *gid10::3xFLAG-GID10 gid5::GID5-3xHA-KANMX* | This study |
| CRLY326 | BY4741; *gid4::3xFLAG-GID4 gid5::Gid5-3xHA-KANMX* | This study |
| CRLY353 | BY4741; *art2::ART2-3xFLAG-HPHNT1* | This study |
| CRLY365 | BY4741; *gid10::3xFLAG-GID10 gid8::GID8-3xHA-KANMX gid4::NATNT2* | This study |
| CRLY382 | BY4741; *lyp1::LYP1-GFP-HPHNT1* | This study |
| CRLY384 | BY4741; *gid4::KANMX lyp1::LYP1-GFP-HPHNT1* | This study |
| CRLY386 | BY4741; *gid10::KANMX lyp1::LYP1-GFP-HPHNT1* | This study |
| CRLY388 | BY4741; *gid2::KANMX lyp1::LYP1-GFP-HPHNT1* | This study |
| CRLY407 | BY4741; *art2::KANMX lyp1::LYP1-GFP-HPHNT1* | This study |
| CRLY431 | BY4741; *art2::KANMX* | This study |
| CRLY434 | BY4741; *art1::NATNT2* | This study |
| CRLY435 | BY4741; *art1::NATNT2 can1::CAN1-GFP-HPHNT1* | This study |
| CRLY451 | BY4741; *art1::NATNT2 lyp1::LYP1-GFP-HPHNT1* | This study |
| CRLY453 | BY4741; *art2::KANMX art1::NATNT2 lyp1::LYP1-GFP-HPHNT1* | This study |
| CRLY458 | BY4741; *art1::ART2(P2S)* | This study |
| CRLY468 | BY4741; *mdh2::MDH2-3xFLAG-HPHNT1* | This study |
| CRLY473 | BY4741; *gid5::KANMX lyp1::LYP1-GFP-HPHNT1* | This study |
| CRLY487 | BY4741; *art2::ART2(P2S)-3xFLAG-HPHNT1* | This study |
| CRLY507 | BY4741; *art2::NATNT2-P<sub>GPD</sub>-ART2 art1::KANMX* | This study |
| CRLY517 | BY4741; *art2::KANMX art1::NATNT2* | This study |
| CRLY569 | BY4741; *gid4::3xFLAG-GID4 gid10::NATNT2-P<sub>GPD</sub>-GID10* | This study |
| LHY146 | BY4741; *gid4::NATNT2 gid10::KANMX* | This study |
| VBY104 | BY4741; *gid4::3xFLAG-GID4 gid5::GID5(W606A, Y613A, Q649A)-3xHA-KANMX* | This study |
| VBY105 | BY4741; *gid10::3xFLAG-GID10 gid5::GID5(W606A, Y613A, Q649A)-3xHA-KANMX* | This study |
| VBY106 | BY4741; *gid4::3xFLAG-GID4(F359A, F361A) gid5::GID5-3xHA-KANMX* | This study |
| VBY107 | BY4741; *gid10::3xFLAG-GID10(Δ289-292) gid5::GID5-3xHA-KANMX* | This study |
| VBY109 | BY4741; *gid4::GID4(F359A, F361A) gid10::3xFLAG-GID10* | This study |
| VBY110 | BY4741; *gid4::3xFLAG-GID4 gid2::KANMX* | This study |
| VBY111 | BY4741; *gid4::3xFLAG-GID4 gid10::KANMX* | This study |
| VBY119 | BY4741; *gid2::KANMX art1::HPHNT1* | This study |
| VBY120 | BY4741; *gid4::KANMX art1::HPHNT1* | This study |
| VBY121 | BY4741; *gid5::KANMX art1::HPHNT1* | This study |
| VBY124 | *BY4741; gid10::KANMX art1::HPHNT1* | This study |
| VBY130 | *BY4741; gid4::KANMX art1::NATNT2 lyp1::LYP1-GFP-HPHNT1* | This study |
| VBY131 | *BY4741; gid10::KANMX art1::NATNT2 lyp1::LYP1-GFP-HPHNT1* | This study |
| VBY132 | *BY4741; gid2::KANMX art1::NATNT2 lyp1::LYP1-GFP-HPHNT1* | This study |
| VBY133 | *BY4741; gid5::KANMX art1::NATNT2 lyp1::LYP1-GFP-HPHNT1* | This study |
| VBY155 | *BY4741; gid4::KANMX gid10::NATNT2 art1::HISMx6 lyp1::LYP1-GFP-HPHNT1* | This study |
| VBY156 | *BY4741; gid4::KANMX art1::NATNT2 can1::CAN1-GFP-HPHNT1* | This study |
| VBY157 | *BY4741; gid10::KANMX art1::NATNT2 can1::CAN1-GFP-HPHNT1* | This study |
| VBY158 | *BY4741; gid2::KANMX art1::NATNT2 can1::CAN1-GFP-HPHNT1* | This study |
| VBY159 | *BY4741; gid5::KANMX art1::NATNT2 can1::CAN1-GFP-HPHNT1* | This study |
| Y2HGOLD | *MATa, trp1-901, leu2-3, 112, ura3-52, his3-200, gal4Δ, gal80Δ, LYS2::GAL1<sub>UAS</sub>-Gal1<sub>TATA</sub>-His3, GAL2<sub>UAS</sub>-Gal2<sub>TATA</sub>-Ade2 URA3::MEL1<sub>UAS</sub>-Mel1<sub>TATA</sub> AUR1-C MEL1* | Takara Bio<br>(Cat#630496) |

### Yeast strains and growth conditions (incl. spot tests)

All yeast strains were constructed using standard techniques [62-64]. Yeast were grown in YPD (1% yeast extract, 2% peptone, 2% glucose) or SD complete (0.67% yeast nitrogen base without amino acids, 2% glucose, containing 87.5 mg/L alanine, arginine, asparagine, aspartic acid, cysteine, glutamine, glutamic acid, glycine, leucine, lysine, methionine, myo-inositol, isoleucine, phenylalanine, proline, serine, threonine, tyrosine and valine, 43.7 mg/L histidine, tryptophan and uracil, 22.5mg/L adenine, and 8.7 mg/L para-aminobenzoic acid) media. Where plasmids are used, the appropriate amino acids were omitted from SD complete media. YPE and SE growth media indicate replacement of the glucose in YPD or SD complete, respectively, with 2% ethanol. For nutrient starvation, yeast cultures were grown to OD600=1.0 in SD complete, washed once with pre-warmed SD-AA (0.67% yeast nitrogen base without amino acids, 2% glucose, and 20mg/L uracil) or SD-N (0.17% yeast nitrogen base without amino acids or ammonium sulfate, 2% glucose), resuspended in pre-warmed SD-AA or SD-N to an OD600=1, and grown for the indicated timepoints. Unless otherwise specified, yeast cultures were grown at 30°C.

### Yeast growth assays

For yeast two-hybrid experiments, pGADCg- and pGBKCg-based plasmids containing the indicated protein fusions were transformed into the yeast strain Y2HGOLD (Takara Bio), and double transformants were selected by growth on SD media lacking leucine and tryptophan. Cells were then grown in SD media lacking leucine and tryptophan and supplemented with 0.1 mg/mL adenine to and OD600 of 1.0-2.0. A 240 μL dilution containing 0.096 ODs of cells was transferred to the first column of a sterile 96-well microtiter plate. Serial dilutions were made, at the dilutions specified, and yeast cells were spotted using a 48 Pin Multi-Blot Replicator (V&P Scientific VP480) on SD media lacking leucine and tryptophan, supplemented with 0.1 mg/mL adenine, and SD media lacking leucine, tryptophan, histidine and adenine (and, where indicated, supplemented with Aureobasidin A). Plates were grown at 30°C.

For growth assays on thialysine, yeast cells were grown in SD complete media to an OD of 1.0-2.0. Yeast cells were diluted as described above and spotted on SD lacking lysine, or SD lacking lysine supplemented with thialysine at the indicated concentrations.

### Yeast cell lysis and western blotting

Protein degradation assays using the promoter reference technique were done as previously described [41]. Cells were transformed with plasmid expressing a test substrate and DHFR from identical promoters containing tetracycline-repressible RNA-binding elements. Yeast cells were grown in SD media lacking histidine to an OD600 of 1.0-1.5, pelleted by centrifugation at 3,000 rpm for 3 minutes, washed once with pre-warmed SE media lacking histidine, resuspended to an OD600 of 1.0 in pre-warmed SE media lacking histidine, and grown for 19 hours. Cells were then pelleted by centrifugation at 3,000 rpm for 3 minutes, resuspended to an OD600 of 1.0 in SD media lacking histidine and allowed to recover for the indicated timepoints. At each timepoint 1 OD of yeast cells was pelleted, supernatant removed, flash frozen in liquid nitrogen and stored at -80°C until lysis. For lysis, yeast cells were resuspended in 0.8 mL 0.2 M NaOH, incubated 20 minutes on ice, and then pelleted by centrifugation at 11,200*xg* for 2 minutes. The supernatant was removed and the pellet was resuspended in 50 μL HU buffer and heated at 70°C for 10 minutes. Lysates were then precleared by centrifugation at 11,200*xg* for 5 minutes and loaded onto a 12% SDS-PAGE gel. Samples were transferred to nitrocellulose membranes and visualized by western blot using αFLAG (Sigma, F1804) and αHA (Sigma H6908) primary antibodies, and Dylight 633 goat anti-Mouse (Invitrogen 35512) and Dylight 488 goat anti-rabbit (Invitrogen 35552) secondary antibodies. Proteins were imaged on an Amersham typhoon scanner (GE Lifesciences), and bands were quantified with ImageStudio software (Licor).

For visualization of 3xFLAG-Gid4, 3xFLAG-Gid10, and Lyp1-GFP (where protein levels are not quantified), cells were grown under the indicated conditions, 5 ODs for 3xFLAG-Gid4 and 3xFLAG-Gid10 or 1.5 ODs for Lyp1-GFP of yeast cells were pelleted at each timepoint, and lysed as described above. Samples were run on a 12% SDS-PAGE gel, transferred to nitrocellulose membrane, and visualized by western blot using αFLAG (Sigma, F1804) or αGFP (Roche 11814460001), and on a separate blot αPGK (Invitrogen 459250) primary antibodies, and goat anti-mouse peroxidase secondary antibody (Sigma A4416). Proteins were visualized on Amersham ImageQuant800 (GE Lifesciences). For visualization of 3xFLAG-Gid4 (where protein levels are quantified), 1.5 OD of yeast cells were pelleted at each timepoint and lysed as described above. Samples were run on a 12% SDS-PAGE gel, transferred to nitrocellulose membrane, and visualized by western blotting using αFLAG (Sigma, F1804) and αPGK (Invitrogen 459250) primary antibodies on the same blot, and Dylight 633 goat anti-Mouse (Invitrogen 35512) secondary antibody.

### Preparation of plasmids for recombinant protein expression

All constructs for bacterial protein expression were prepared by Gibson assembly method [65]. For generation of mutant versions of the genes, the QuickChange mutagenesis protocol was applied (Stratagene). All coding sequences used for protein expression were verified by DNA sequencing. To express the GID complex in insect cells from a single baculoviral expression vector, genes encoding GID subunits were combined by the biGBac assembly method [66].

### Recombinant protein expression and purification

Both WT and mutant versions of the GID complex used for biochemical assays were expressed in Hi-5 insect cells transfected with recombinant baculovirus variants in EX-CELL 420 Serum-Free Medium. After 72 hours at 27°C, the cultures were harvested and resuspended in a lysis buffer containing 50 mM HEPES pH 7.5, 200 mM NaCl, 5 mM βDTT, 10 μg/ml leupeptin, 20 μg/ml aprotinin, 2 mM benzamidine, EDTA-free cOmplete protease inhibitor tablet (Roche, 1 tablet per 50 ml of buffer) and 1 mM PMSF. The complex was first affinity purified via a twin-Strep tag appended to Gid8 C-terminus. Further purification was performed by anion exchange chromatography and size exclusion chromatography (SEC) in the final buffer containing 25 mM HEPES pH 7.5, 200 mM NaCl and 1 mM DTT.

Aside from the GID complex, all recombinant proteins were expressed in *E. coli* BL21 (DE3) RIL. Cells transformed with an appropriate expression plasmid were grown in Terrific Broth (TB) medium at 37°C until OD600 of 0.6 and cooled down to 18°C. Then, overnight expression of proteins was induced by addition of 0.4 mM IPTG. All versions of Gid4 and Gid10 were expressed as GST-TEV fusions. After harvesting, cell pellets were resuspended in the lysis buffer containing 50 mM HEPES pH 7.5, 200 mM NaCl, 5 mM DTT and 1 mM PMSF. GST-tagged proteins were purified from bacterial lysates by glutathione affinity chromatography, followed by overnight digestion at 4°C with tobacco etch virus (TEV) protease to cleave off the GST tag. Further purification was carried out with SEC in the buffer containing 25 mM HEPES pH 7.5, 150 mM NaCl and 1 mM DTT, 5 mM DTT or 0.5 mM TCEP for biochemical assays, crystallography and ITC binding test, respectively. At the end, a pass-back over glutathione affinity resin was performed to get rid of the remaining uncleaved GST-fusion protein and free GST. Ubc8 was expressed with a C-terminal 6xHis-tag. After harvesting, cell pellet was resuspended in the lysis buffer containing 50 mM HEPES pH 7.5, 200 mM NaCl, 5 mM Δ-mercaptoethanol, 10 mM imidazole and 1 mM PMSF. Ubc8-6xHis was purified by nickel affinity chromatography, followed by anion exchange and SEC. Untagged WT ubiquitin was purified via glacial acetic acid method (Kaiser et al., 2011), followed by gravity S column ion exchange chromatography and size exclusion chromatography.

### Immunoprecipitation and ubiquitination assay

For Art2 IPs, yeast cells were grown in YPD at 30°C to an OD600 of 1.0-2.0. For Mdh2 IPs, yeast cells were grown in YPD to an OD600 of 1.0-1.5, pelleted by centrifugation at 3,000 rpm for 3 minutes, washed once with pre-warmed YPE, resuspended to an OD600 of 1.0 in fresh, pre-warmed YPE, and grown at 30°C for 19 hours. 100 ODs of cells were pelleted by centrifugation at 3,000 rpm for 3 min, washed with dH20, resuspended in 1 mL lysis buffer (50 mM Tris-HCL, pH 7.5, 150 mM NaCl, 2 mM EDTA, 50 mM NaF, 0.1% SDS, 1% NP-40, 0.5% Na-deoxycholate, 20 mM NEM, 1% glycerol, and cOmplete EDTA-free protease inhibitor tablets (Roche)), and transferred to a 2 mL tube containing lysing matrix C (MP Biomedicals). Cells were lysed by 3 rounds of 20 seconds in a Fast-Prep24 instrument (MP Biomedicals), resting on ice for 5 minutes in between each round. Lysates were then pre-cleared by centrifugation at 4,000*xg* for 10 minutes. The supernatant was added to 50 μL pre-washed anti-DYKDDDDK magnetic agarose beads (ThermoFischer A36797) and nutated at 4°C for 2 hours. Beads were then pelleted on a magnetic rack, and supernatant was discarded. Beads were washed twice with wash buffer 1 (50 mM Tris-HCL, pH 7.5, 150 mM NaCl, 2 mM EDTA, 50 mM NaF, 0.1% SDS, 1%NP-40, 0.5% Na-deoxycholate, 20 mM NEM, 1% glycerol), twice with wash buffer 2 (25 mM HEPES, pH 7.5, 100 mM NaCl), and resuspended in 100 μL of wash buffer 2. For each ubiquitination reaction, 25 μL of this suspension were pelleted on a magnetic rack, and supernatant removed. Ubiquitination reaction mix (1 μM E2 Ubc8-6xHis, 0.5 μM GID^Ant^, 30 μM Ubiquitin, 0.5 μM substrate receptor (none, Gid10^57-292^, Gid10^57-288^, or Gid4^117-362^, as indicated), 25 mM HEPES, pH 7.5, 100 mM NaCl, 2.5 mM MgCl2, and 1 mM ATP) was added to the beads. The reaction was started by addition of 0.2 μM E1 Uba1 and incubated at room temperature for the indicated timepoints. Beads were then pelleted on a magnetic rack, washed with wash buffer 1, resuspended in 30 μL 2x sample buffer, and heated at 95°C for five minutes to elute the protein. For Art2-3xFLAG blots, the eluate was loaded on 4-12% SDS-PAGE gels, run at 200V for 80 minutes, and transferred to PVDF membrane at 100V for 90 minutes. For Mdh2-3xFLAG blots, eluate was loaded on a 12% SDS-PAGE gel, run at 200V for 50 minutes, and transferred to nitrocellulose membrane at 100V for 60 minutes. Samples were then visualized by immunoblotting with anti-FLAG (Sigma F1804) primary antibody and goat anti-mouse peroxidase secondary antibody (Sigma A4416), and imaged on an Amersham ImageQuant800 (GE Lifesciences).

### *In vitro* binding assay

To test if the GID complex binds Gid10 in a manner similar to Gid4, WT and mutant versions (Gid5 W606A/Y613A/Q649A) of GID^Ant^ were mixed with two-fold molar excess of Gid10^57-292^, Gid10^57-288^, Gid4^117-358^ and Gid4^117-362^. After incubating the proteins for 30 minutes on ice, 20 µL of Strep-Tactin resin was added to the mixture and further incubated for 30 minutes. After thorough wash of the resin, proteins were eluted and analyzed with SDS-PAGE.

### Isothermal titration calorimetry (ITC)

The ITC measurements were carried out with MicroCal PEAQ-ITC instrument (Malvern Panalytical) at 25°C. The Art2 degron peptides were dissolved in the Gid10 SEC buffer containing 25 mM HEPES pH 7.5, 150 mM NaCl and 0.5 mM TCEP and their concentration was measured by absorbance at 280 nm. Binding experiments were carried out by titrating 198 or 450 µM peptides to Gid10^57-292^ at 21 or 42 µM for PFITSRPW and PFITSRPVAW, respectively. Peptides were added to Gid10 by nineteen 2 µl injections, with 4 s injection time and 150 s equilibration between the injections. The reference power was set to 10 µcal/s. Raw ITC data were analyzed using one site binding mode in MicroCal ITC analysis software (Malvern Panalytical) to determine *K*D and stoichiometry of the binding reaction. All plots were prepared in GraphPad Prism.

### *In vitro* ubiquitylation assay

To verify whether Art2 N-terminus can be ubiquitinated by GID^SR10^, we performed an *in vitro* activity assay with Art2^2-28^ WT and P2S mutant peptides, with fluorescein appended to their C-termini. Ubiquitination reaction was performed in a multi-turnover format in a buffer containing 25 mM HEPES pH 7.5, 150 mM NaCl, 5 mM ATP and 10 mM MgCl2. To start the reaction, 0.2 µM E1 Uba1, 1 µM E2 Ubc8-6xHis, 0.5 µM E3 GID^Ant^, 20 μM Ub, 0 or 1 µM Gid4^117-362^ or Gid10^57-292^ and 1 µM peptide substrate were mixed and incubated at room temperature. At indicated timepoints, an aliquot of the reaction mix was mixed with SDS-PAGE loading buffer. The outcome of the activity assay was visualized with a fluorescent scan of an SDS-PAGE gel with Typhoon imager (GE Healthcare).

### Gid10 crystallization, data collection and structure determination

Crystallization trials were carried out in the MPIB crystallography facility. Before setting up crystallization trays, Gid10^65-284^ was concentrated and mixed with PFITSRPW peptide to obtain final concentration of protein and peptide of 262 μM and 760 μM, respectively (∼3-fold molar excess of the peptide). The crystal that gave rise to the final structure was grown at room temperature in the buffer containing 18.5% PEG3350, 0.1M Bis-Tris propane pH 6.0 and 0.2M potassium chloride by vapor diffusion in a sitting-drop format. Before data collection, crystals were cryoprotected in 20% ethylene glycol and flash-frozen in liquid nitrogen.

Diffraction dataset was recorded at X10SA beam line, Swiss Light Source (SLS) in Villingen, Switzerland. Data were recorded at 0.5-degree rotation intervals using Dectris Eiger II 16 M detector. Data were indexed, integrated, and scaled using XDS package to a resolution limit of 1.3 Å. Phasing was performed through molecular replacement using a structure of yeast Gid4 (extracted from PDB: 7NS3) with PHASER integrated into the PHENIX software suite [67-69]. Model building was done using Coot [70,71], whereas refinement was carried out with phenix.refine. Details of X-ray diffraction data collection and refinement statistics are listed in Table S1.

### Proteomics sample preparation

Samples were prepared and analyzed as previously described [40]. Briefly, sodium deoxycholate (SDC) lysis buffer (1% SDC and 100 mM Tris, pH 8.5) were added to the frozen cell pellets. Lysates were immediately boiled for 5 min at 95 °C and homogenized with sonication. Protein concentrations were estimated by tryptophan assay. Equal protein amounts were reduced and alkylated using CAA and TCEP, final concentrations of 40 mM and 10 mM, respectively, for 5 min at 45 °C. Samples were digested overnight at 37°C using trypsin (1:100 wt/wt; Sigma-Aldrich) and LysC (1/100 wt/wt; Wako). Next, peptides were desalted using SDB-RPS StageTips (Empore). Samples were first diluted with 1% trifluoroacetic acid (TFA) in isopropanol to a final volume of 200μL and loaded onto StageTips and subsequently washed with 200μL of 1% TFA in isopropanol twice and 200μL of 0.2% TFA/2% ACN (acetonitrile). Peptides were eluted with 80μl of 1.25% Ammonium hydroxide (NH4OH)/80% ACN, dried using a SpeedVac centrifuge (Concentrator Plus; Eppendorf) and resuspended in buffer A* (0.2% TFA/2% ACN) prior to LC-MS/MS analysis. Peptide concentrations were measured optically at 280 nm (Nanodrop 2000; Thermo Scientific) and subsequently equalized using buffer A*. Three hundred nanograms of peptide was subjected to LC-MS/MS analysis.

### LC-MS/MS Measurements

Samples were loaded onto a 20-cm reversed-phase column (75-μm inner diameter, packed in-house with ReproSil-Pur C18-AQ1.9 μm resin [Dr. Maisch GmbH]). The column temperature was maintained at 60 °C using a homemade column oven. A binary buffer system, consisting of buffer A (0.1% formic acid [FA]) and buffer B (0.1% FA and 80% ACN), was used for peptide separation, at a flow rate of 450 nL/min. An EASY-nLC1200 system (Thermo Fisher Scientific), coupled with the mass spectrometer (Q Exactive HF-X, Thermo Fisher Scientific) via a nano-electrospray source, was employed for nano-flow liquid chromatography. We used a gradient starting at 5% buffer B, increased to 35% in 18.5 min 95% in a minute, and stayed at 95% for min. The mass spectrometer was operated in data independent acquisition mode (DIA). Full MS resolution was set to 120,000 with a full scan range of 300 to 1,650 m/z, a maximum fill time of 60 ms, and an AGC target of 3e6. One full scan was followed by 12 windows with a resolution of 30,000 in profile mode. Precursor ions were fragmented by stepped HCD (NCE 25.5, 27, and 30%).

### Data Processing and Bioinformatics Analysis

DIA files were analyzed using the proteome library previously generated [40] with default settings and enabled cross-run normalization using Spectronaut version 13 (Biognosys). The Perseus software package versions 1.6.0.7 and 1.6.0.9 [72] and GraphPad Prism version 7.03 were used for the data analysis. Protein intensities were log2-transformed and the datasets were filtered to make sure that identified proteins showed expression or intensity in all biological triplicates of at least one condition and the missing values were subsequently replaced by random numbers that were drawn from a normal distribution (width=0.3 and downshift=1.8) in Perseus. To determine significantly different proteins, two sample t-test was applied, assuming that variance within the groups of replicates was equal.

## Acknowledgements

We thank R. Prabu und J. Basquin for assistance with crystallography; A. Varshavsky for providing PRT and Y2H plasmids; I. Paron for technical assistance; the Paul Scherrer Institut (Villigen, Switzerland) for provision of synchrotron radiation beamtime at beamline X10SA of the SLS; A. C. Michaelis, L. Hehl, F. Wilfling, S. Gronau and all members of the Departments of Molecular Machines and Signaling and Proteomics and Signal Transduction for their advice and support.

This study was supported by the Max Planck Gesellschaft, the European Research Council (ERC) under the European Union’s Horizon 2020 research and innovation programme (grant agreement No 789016-NEDD8Activate), and the Gottfried Wilhelm Leibniz Prize from the Deutsche Forschungsgemeinschaft (DFG) (grant SCHU 3196/1-1).

## Author Contributions

Conceptualization: CRL and BAS; Methodology and Investigation: CRL, VB, OK, DS, and JC; Writing-Original Draft: CRL; Writing-Review & Editing: CRL, VB, OK, DS, JC, and BAS; Supervision: CRL, MM, and BAS; Funding Acquisition: MM and BAS.

The authors declare that they have no conflict of interest.

---

**Figure EV1.**
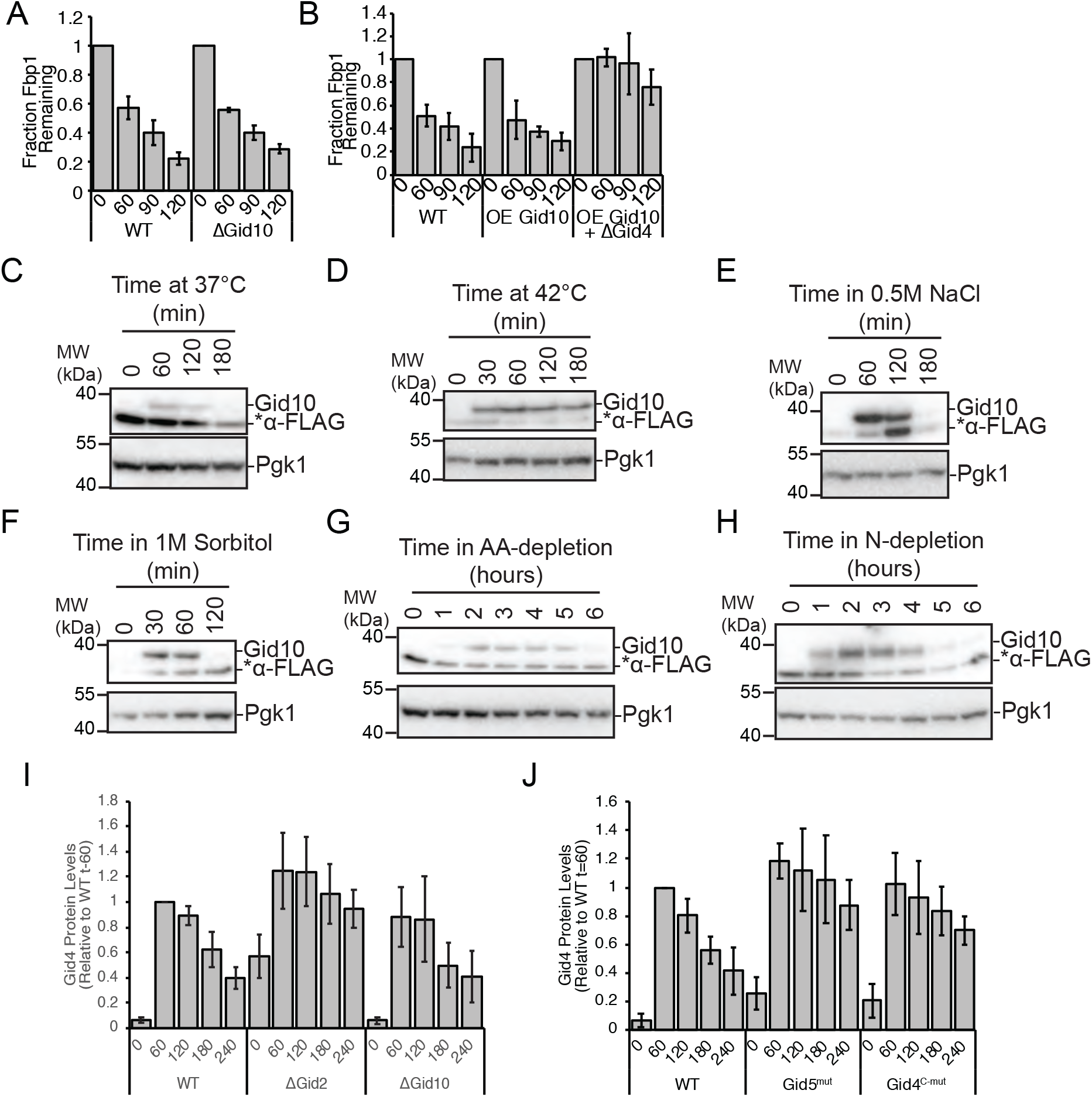
Regulation of Gid10 and Gid4 expression. A) Tetracycline reference-based chase performed during transition from ethanol to glucose media with wildtype and ΔGid10 strains. Bars represent mean, error bars represent standard deviation (n>3). B) Tetracycline reference-based chase performed during transition from ethanol to glucose media with wildtype, and wildtype and ΔGid4 strains overexpressing (OE) Gid10. Bars represent mean, error bars represent standard deviation (n>3). C) Lysates from a yeast strain expressing endogenously tagged 3xFLAG-Gid10 that was grown in YPD at 37°C for the indicated timepoints were run on an SDS-PAGE gel and immunoblotted with αFLAG and αPGK. D) Lysates from a yeast strain expressing endogenously tagged 3xFLAG-Gid10 that was grown in YPD at 42°C for the indicated timepoints were run on an SDS-PAGE gel and immunoblotted with αFLAG and αPGK. E) Lysates from a yeast strain expressing endogenously tagged 3xFLAG-Gid10 that was grown in YPD supplemented with 0.5M NaCl for the indicated timepoints were run on an SDS-PAGE gel and immunoblotted with αFLAG and αPGK. F) Lysates from a yeast strain expressing endogenously tagged 3xFLAG-Gid10 that was grown in SD complete supplemented with 1M Sorbitol for the indicated timepoints were run on an SDS-PAGE gel and immunoblotted with αFLAG and αPGK. G) Lysates from a yeast strain expressing endogenously tagged 3xFLAG-Gid10 that was grown in SD-AA for the indicated timepoints were run on an SDS-PAGE gel and immunoblotted with αFLAG and αPGK. H) Lysates from a yeast strain expressing endogenously tagged 3xFLAG-Gid10 that was grown in SD-N for the indicated timepoints were run on an SDS-PAGE gel and immunoblotted with αFLAG and αPGK. I) Wildtype, ΔGid2 and ΔGid10 yeast strains expressing endogenously tagged 3xFLAG-Gid4 were grown in YPE for 19 hours and then shifted to YPD for the indicated timepoints. Lysates were run on an SDS-PAGE gel and immunoblotted with αFLAG and αPGK (as a reference control). Bars represent mean, error bars represent standard deviation (n>3). J) Wildtype, Gid5^W606A, Y613A, Q649A^, and Gid4^F359A,F361A^ (c-mut) yeast strains expressing endogenously tagged 3xFLAG-Gid4 were grown in YPE for 19 hours and then shifted to YPD for the indicated timepoints. Lysates were run on an SDS-PAGE gel and immunoblotted with αFLAG and αPGK (as a reference control). Bars represent mean, error bars represent standard deviation (n>3).

**Figure EV2.**
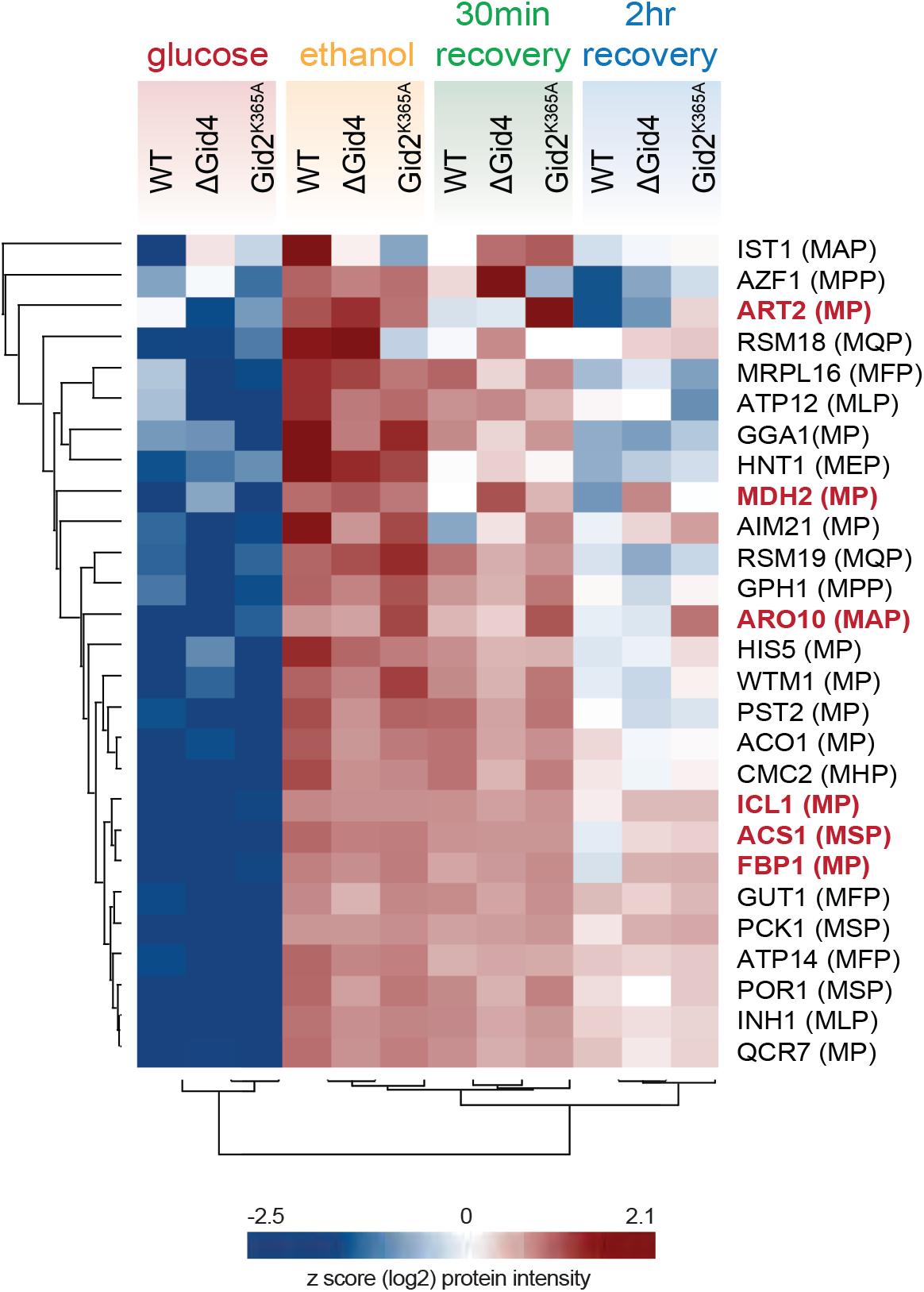
Protein expression during recovery from ethanol starvation. Heat map of z-scored abundances (log2) of the proteins which have the following criteria: 1) significantly upregulated in ethanol compared to glucose, 2) significantly upregulated in ethanol compared to 2 hour recovery, and 3) contains a proline in position 2 or 3.

**Figure EV3.**
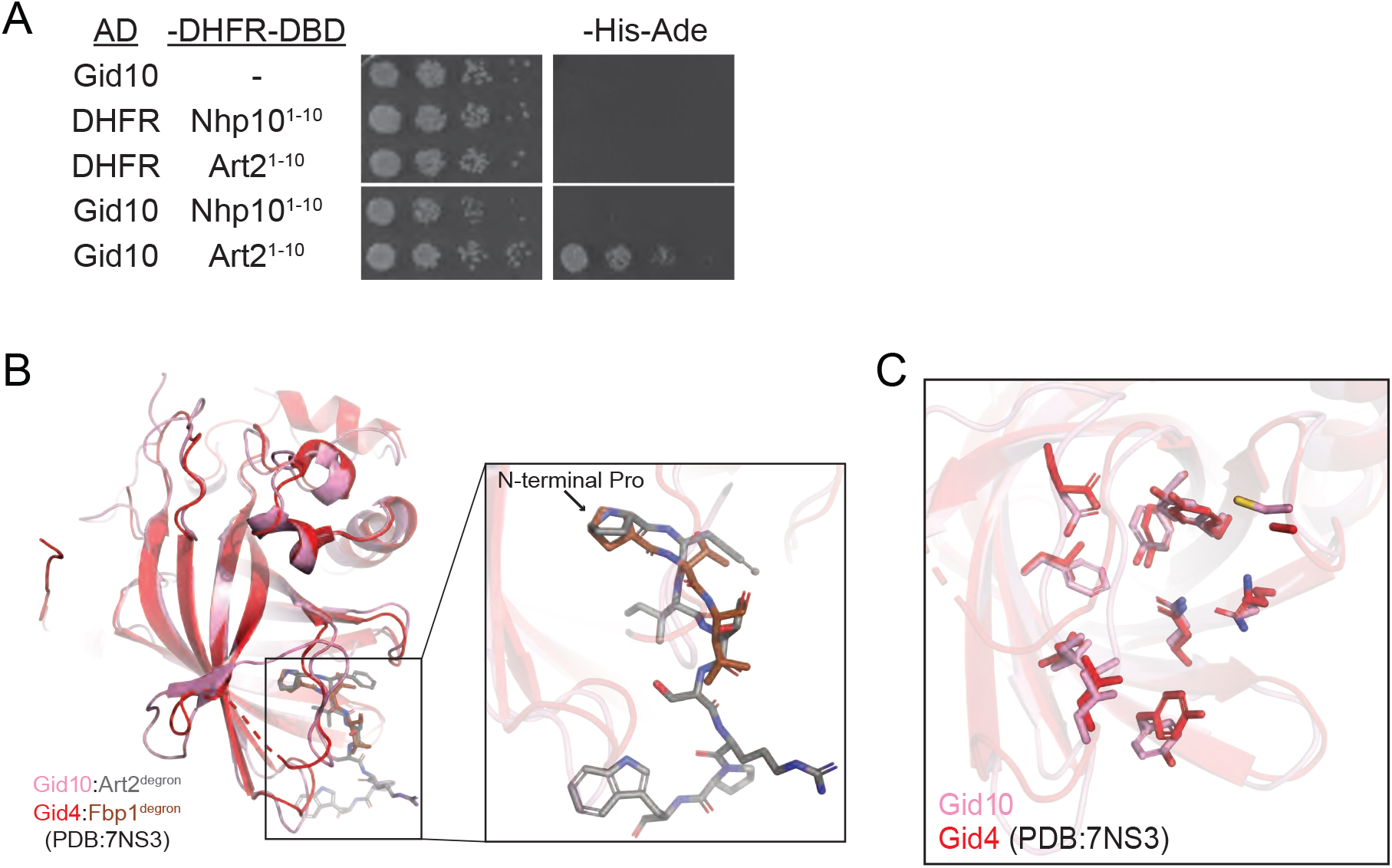
Gid10 interacts with the Art2 N-terminus. A) Yeast two-hybrid between SR-Gal4 activation domain (AD) and substrate degrons fused to DHFR-DNA binding domain (-DHFR-DBD). Growth on -His-Ade is indicative of an interaction between the two test proteins. Spots represent 1:5 serial dilutions. B) Overlay of Gid10 (pink): Art2^2-8^ (grey) and Gid4 (red): Fbp1^2-4^ (brown) (extracted from PDB:7NS3) showing overall similarity of their substrate binding domains as well as the trajectory of the bound degrons. C) Overlay of Gid10 (pink) and Gid4 (red, extracted from PDB:7NS3) highlighting the residues inside their substrate-binding pockets (shown as sticks).

**Figure EV4.**
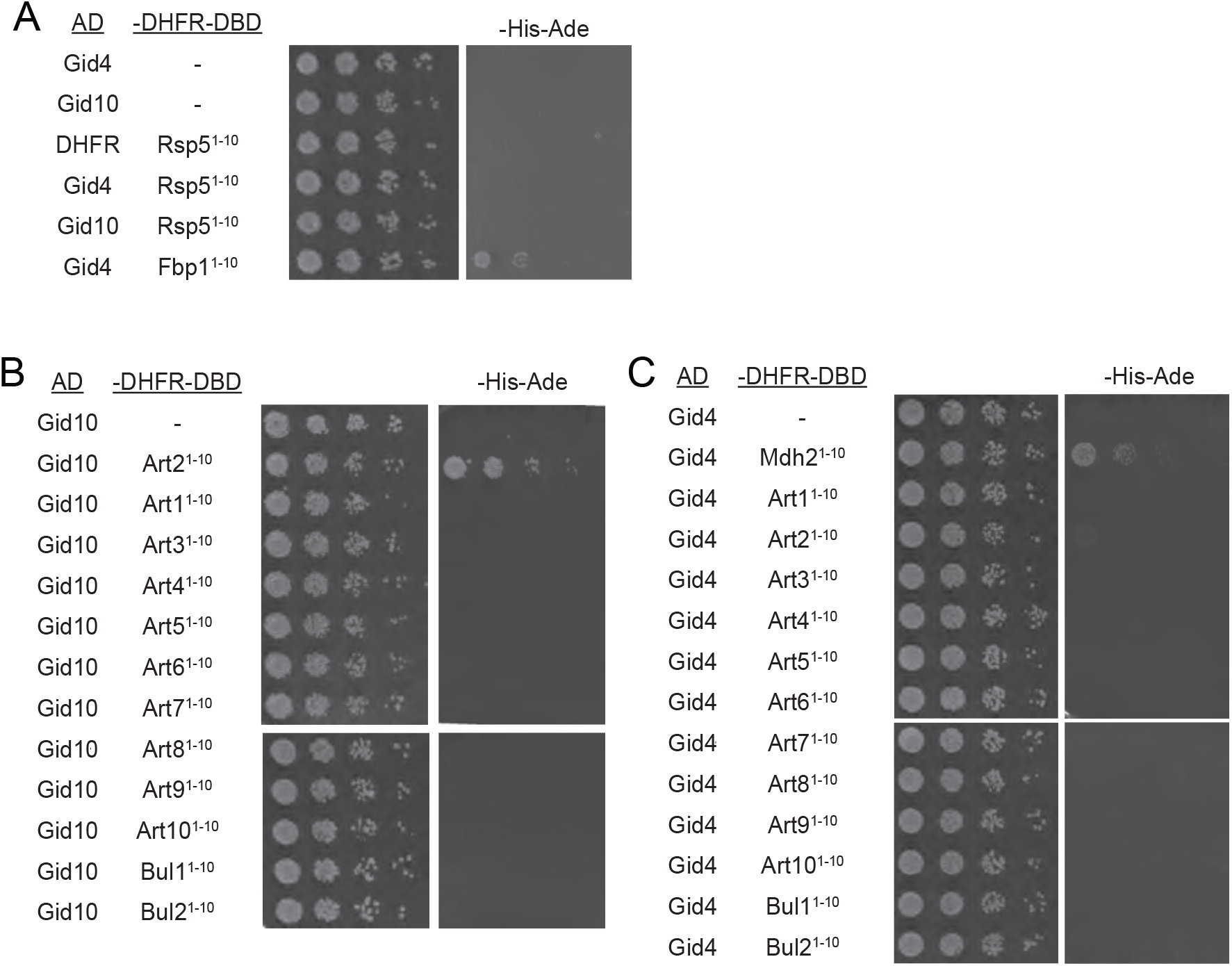
Gid10 and Gid4 do not interact broadly with arrestin degrons. A) Yeast two-hybrid between SR-Gal4 activation domain (AD) and Rsp5 degrons fused to DHFR-DNA binding domain (-DHFR-DBD). Interaction between Gid4-AD and Fbp1-DBD is shown as a control. Growth on -His-Ade is indicative of an interaction between the two test proteins. Spots represent 1:5 serial dilutions. B) Yeast two-hybrid between Gid10-Gal4 activation domain (AD) and arrestin degrons fused to DHFR-DNA binding domain (-DHFR-DBD). Growth on -His-Ade is indicative of an interaction between the two test proteins. Spots represent 1:5 serial dilutions. C) Yeast two-hybrid between Gid4-Gal4 activation domain (AD) and arrestin degrons fused to DHFR-DNA binding domain (-DHFR-DBD). Interaction between Gid4-AD and Mdh2-DBD is shown as a control. Growth on -His-Ade is indicative of an interaction between the two test proteins. Spots represent 1:5 serial dilutions.

**Figure EV5.**
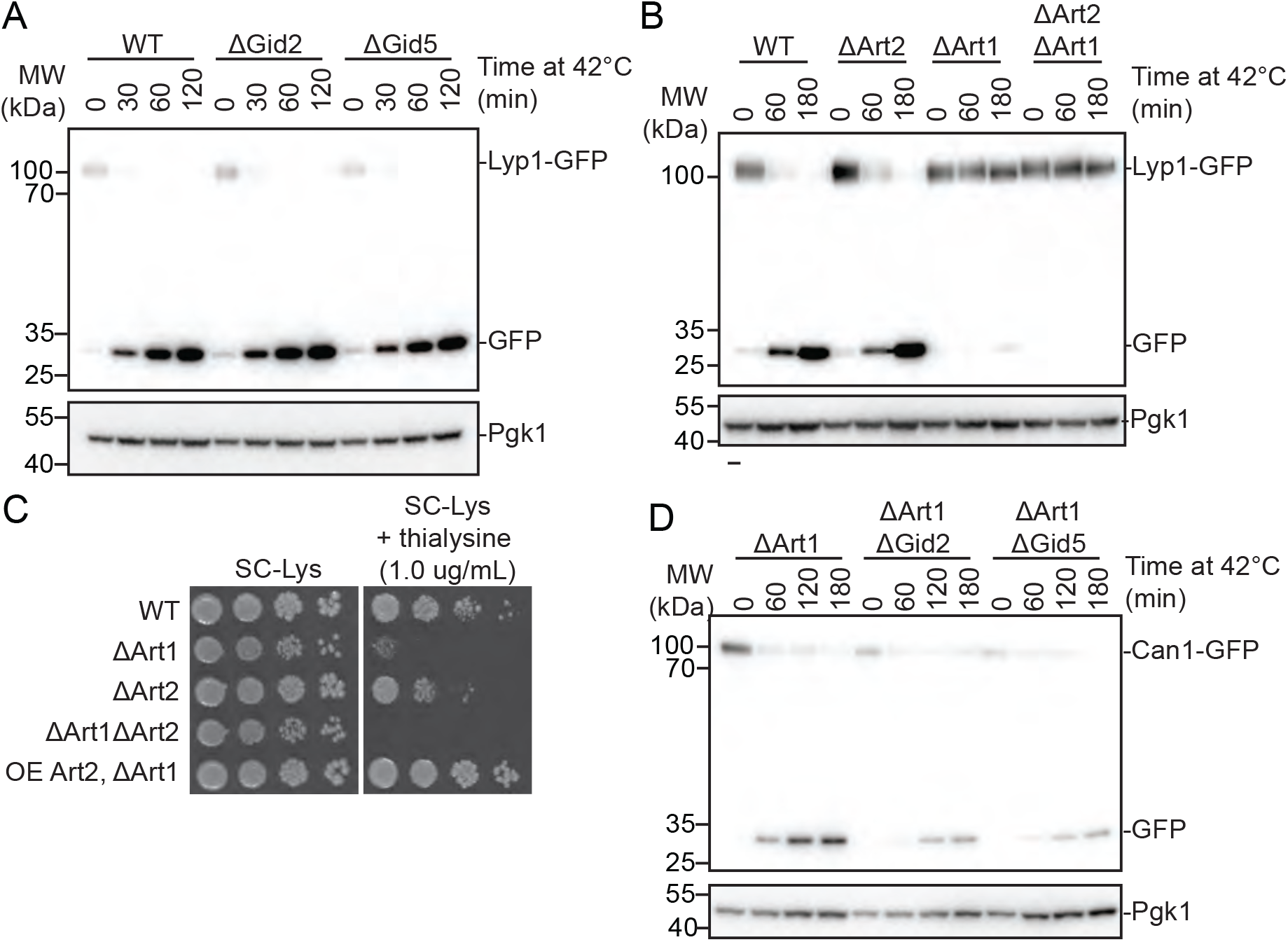
Regulation of amino acid receptors during heat shock. A) Wildtype, ΔGid2 and ΔGid5 yeast strains expressing endogenously tagged Lyp1-GFP were grown at 42°C for the indicated timepoints. Lysates were run on an SDS-PAGE B) Wildtype, ΔArt2, ΔArt1, and ΔArt2ΔArt1 yeast strains expressing endogenously taggedLyp1-GFP were grown at 42°C for the indicated timepoints. Lysates were run on an SDS-PAGE gel and immunoblotted with αGFP and αPGK. C) Growth assay of wildtype, ΔArt2, ΔArt1, ΔArt2ΔArt1, and ΔArt1 overexpressing Art2 yeast strains on SD-Lys (-) and SD-Lys containing 1.0 μg/mL thialysine (+). Spots represent 1:5 serial dilutions. D) ΔArt1 strains containing *GID2* or *GID5* deletions and expressing endogenously tagged Can1-GFP were grown at 42°C for the indicated timepoints. Lysates were immunoblotted for αGFP and αPGK.

**Table S1:**
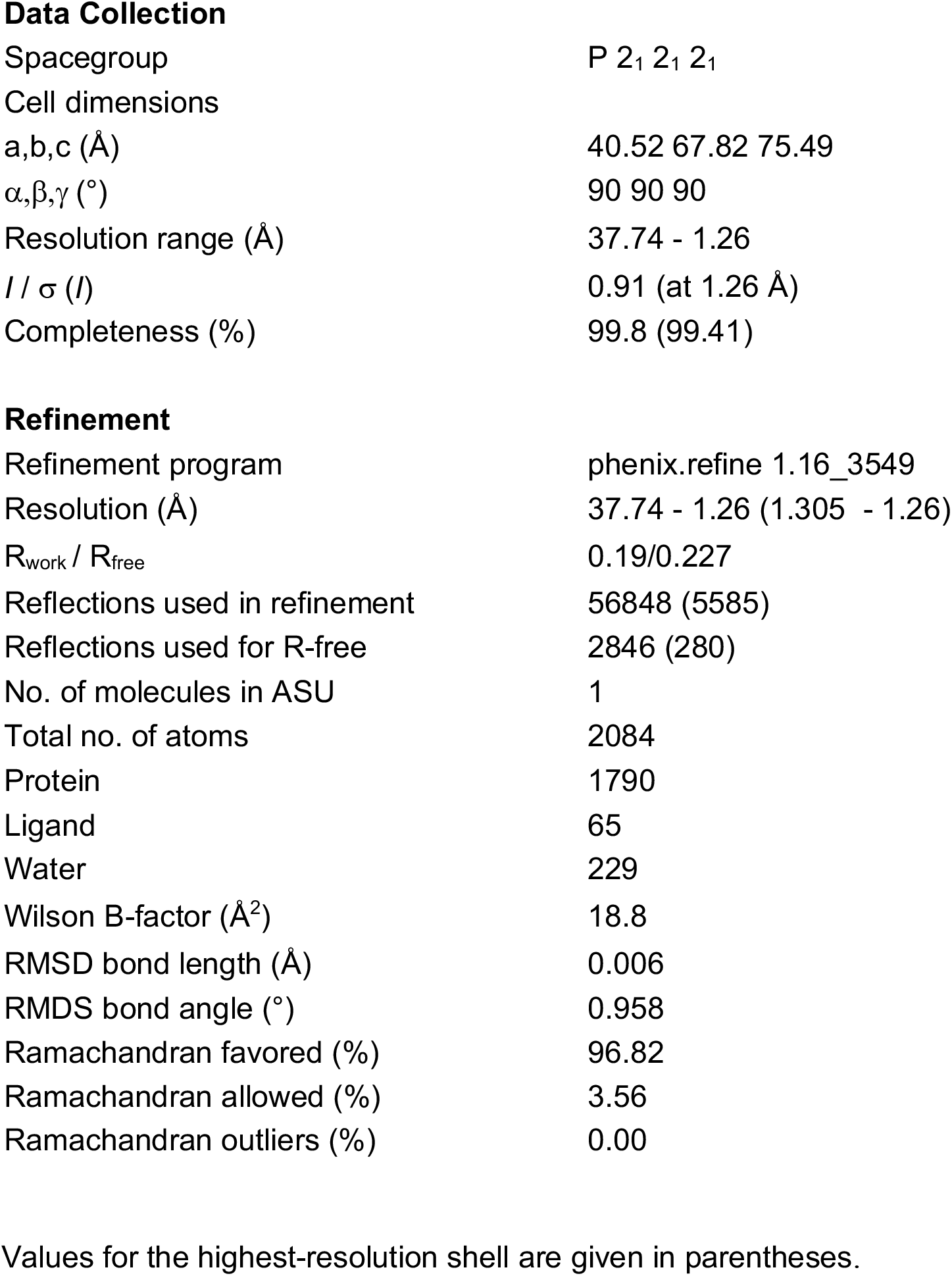
Crystallography data collection and refinement statistics Values for the highest-resolution shell are given in parentheses.

|  |  |
| --- | --- |
| Spacegroup | P 2<sub>1</sub> 2<sub>1</sub> 2<sub>1</sub> |
| Cell dimensions |  |
| a,b,c (Å) | 40.52 67.82 75.49 |
| $\alpha,\beta,\gamma$ (°) | 90 90 90 |
| Resolution range (Å) | 37.74 - 1.26 |
| $I / \sigma(I)$ | 0.91 (at 1.26 Å) |
| Completeness (%) | 99.8 (99.41) |
| **Refinement** |  |
| Refinement program | phenix.refine 1.16_3549 |
| Resolution (Å) | 37.74 - 1.26 (1.305 - 1.26) |
| R<sub>work</sub> / R<sub>free</sub> | 0.19/0.227 |
| Reflections used in refinement | 56848 (5585) |
| Reflections used for R-free | 2846 (280) |
| No. of molecules in ASU | 1 |
| Total no. of atoms | 2084 |
| Protein | 1790 |
| Ligand | 65 |
| Water | 229 |
| Wilson B-factor (Å<sup>2</sup>) | 18.8 |
| RMSD bond length (Å) | 0.006 |
| RMSD bond angle (°) | 0.958 |
| Ramachandran favored (%) | 96.82 |
| Ramachandran allowed (%) | 3.56 |
| Ramachandran outliers (%) | 0.00 |

## References

1. Cappadocia L, Lima CD (2018) Ubiquitin-like Protein Conjugation: Structures, Chemistry, and Mechanism. Chem Rev 118: 889–918.

2. Deshaies RJ, Joazeiro CAP (2009) RING domain E3 ubiquitin ligases. Annu Rev Biochem 78: 399–434.

3. Rotin D, Kumar S (2009) Physiological functions of the HECT family of ubiquitin ligases. Nat Rev Mol Cell Biol 10: 398–409.

4. Jackson PK, Eldridge AG, Freed E, Furstenthal L, Hsu JY, Kaiser BK, Reimann JD (2000) The lore of the RINGs: substrate recognition and catalysis by ubiquitin ligases. Trends Cell Biol 10: 429–439.

5. Skowyra D, Craig KL, Tyers M, Elledge SJ, Harper JW (1997) F-box proteins are receptors that recruit phosphorylated substrates to the SCF ubiquitin-ligase complex. Cell 91: 209–219.

6. Bai C, Sen P, Hofmann K, Ma L, Goebl M, Harper JW, Elledge SJ (1996) SKP1 connects cell cycle regulators to the ubiquitin proteolysis machinery through a novel motif, the F-box. Cell 86: 263–274.

7. Zheng N, Schulman BA, Song L, Miller JJ, Jeffrey PD, Wang P, Chu C, Koepp DM, Elledge SJ, Pagano M, et al. (2002) Structure of the Cul1-Rbx1-Skp1-F boxSkp2 SCF ubiquitin ligase complex. Nature 416: 703–709.

8. Rusnac D-V, Zheng N (2020) Structural Biology of CRL Ubiquitin Ligases. Adv Exp Med Biol 1217: 9–31.

9. Dupré S, Urban-Grimal D, Haguenauer-Tsapis R (2004) Ubiquitin and endocytic internalization in yeast and animal cells. Biochim Biophys Acta 1695: 89–111.

10. Lin CH, MacGurn JA, Chu T, Stefan CJ, Emr SD (2008) Arrestin-Related Ubiquitin-Ligase Adaptors Regulate Endocytosis and Protein Turnover at the Cell Surface. Cell 135: 714–725.

11. Foot NJ, Dalton HE, Shearwin-Whyatt LM, Dorstyn L, Tan S-S, Yang B, Kumar S (2008) Regulation of the divalent metal ion transporter DMT1 and iron homeostasis by a ubiquitin-dependent mechanism involving Ndfips and WWP2. Blood 112: 4268–4275.

12. Trimpert C, Wesche D, de Groot T, Pimentel Rodriguez MM, Wong V, van den Berg DTM, Cheval L, Ariza CA, Doucet A, Stagljar I, et al. (2017) NDFIP allows NEDD4/NEDD4L-induced AQP2 ubiquitination and degradation. PLoS ONE 12: e0183774.

13. Mund T, Pelham HRB (2009) Control of the activity of WW-HECT domain E3 ubiquitin ligases by NDFIP proteins. EMBO Rep 10: 501–507.

14. Lu PJ, Zhou XZ, Shen M, Lu KP (1999) Function of WW domains as phosphoserine-or phosphothreonine-binding modules. Science 283: 1325–1328.

15. Ogunjimi AA, Briant DJ, Pece-Barbara N, Le Roy C, Di Guglielmo GM, Kavsak P, Rasmussen RK, Seet BT, Sicheri F, Wrana JL (2005) Regulation of Smurf2 ubiquitin ligase activity by anchoring the E2 to the HECT domain. Mol Cell 19: 297–308.

16. Huibregtse JM, Scheffner M, Howley PM (1993) Localization of the E6-AP regions that direct human papillomavirus E6 binding, association with p53, and ubiquitination of associated proteins. Mol Cell Biol 13: 4918–4927.

17. Persaud A, Alberts P, Mari S, Tong J, Murchie R, Maspero E, Safi F, Moran MF, Polo S, Rotin D (2014) Tyrosine phosphorylation of NEDD4 activates its ubiquitin ligase activity. Sci Signal 7: ra95–ra95.

18. Polo S, Di Fiore PP (2008) Finding the right partner: science or ART? Cell 135: 590–592.

19. Shearwin-Whyatt L, Dalton HE, Foot N, Kumar S (2006) Regulation of functional diversity within the Nedd4 family by accessory and adaptor proteins. Bioessays 28: 617–628.

20. Hettema EH, Valdez-Taubas J, Pelham HRB (2004) Bsd2 binds the ubiquitin ligase Rsp5 and mediates the ubiquitination of transmembrane proteins. EMBO J 23: 1279–1288.

21. Plant PJ, Lafont F, Lecat S, Verkade P, Simons K, Rotin D (2000) Apical membrane targeting of Nedd4 is mediated by an association of its C2 domain with annexin XIIIb. The Journal of Cell Biology 149: 1473–1484.

22. Lauwers E, Erpapazoglou Z, Haguenauer-Tsapis R, André B (2010) The ubiquitin code of yeast permease trafficking. Trends Cell Biol 20: 196–204.

23. Zhao Y, MacGurn JA, Liu M, Emr S (2013) The ART-Rsp5 ubiquitin ligase network comprises a plasma membrane quality control system that protects yeast cells from proteotoxic stress. Elife 2: e00459.

24. Becuwe M, Herrador A, Haguenauer-Tsapis R, Vincent O, Léon S (2012) Ubiquitin-Mediated Regulation of Endocytosis by Proteins of the Arrestin Family. Biochemistry Research International 2012: 1–12.

25. Kahlhofer J, Léon S, Teis D, Schmidt O (2020) The α-arrestin family of ubiquitin ligase adaptors links metabolism with selective endocytosis. Biol Cell boc.202000137.

26. Ivashov V, Zimmer J, Schwabl S, Kahlhofer J, Weys S, Gstir R, Jakschitz T, Kremser L, Bonn GK, Lindner H, et al. (2020) Complementary α-arrestin-ubiquitin ligase complexes control nutrient transporter endocytosis in response to amino acids. Elife 9: 1389.

27. French ME, Kretzmann BR, Hicke L (2009) Regulation of the RSP5 ubiquitin ligase by an intrinsic ubiquitin-binding site. J Biol Chem 284: 12071–12079.

28. MacDonald C, Shields SB, Williams CA, Winistorfer S, Piper RC (2020) A Cycle of Ubiquitination Regulates Adaptor Function of the Nedd4-Family Ubiquitin Ligase Rsp5. Curr Biol 30: 465–479.e465.

29. Barnett JA, Entian K-D (2005) A history of research on yeasts 9: regulation of sugar metabolism. Yeast 22: 835–894.

30. Gancedo JM (1998) Yeast carbon catabolite repression. Microbiol Mol Biol Rev 62: 334–361.

31. Hovsepian J, Defenouillère Q, Albanèse V, Váchová L, Garcia C, Palková Z, Léon S (2017) Multilevel regulation of an α-arrestin by glucose depletion controls hexose transporter endocytosis. The Journal of Cell Biology 216: 1811–1831.

32. Hämmerle M, Bauer J, Rose M, Szallies A, Thumm M, Düsterhus S, Mecke D, Entian KD, Wolf DH (1998) Proteins of newly isolated mutants and the amino-terminal proline are essential for ubiquitin-proteasome-catalyzed catabolite degradation of fructose-1,6-bisphosphatase of Saccharomyces cerevisiae. J Biol Chem 273: 25000–25005.

33. Chen S-J, Wu X, Wadas B, Oh J-H, Varshavsky A (2017) An N-end rule pathway that recognizes proline and destroys gluconeogenic enzymes. Science 355: eaal3655–eaal3668.

34. Santt O, Pfirrmann T, Braun B, Juretschke J, Kimmig P, Scheel H, Hofmann K, Thumm M, Wolf DH (2008) The yeast GID complex, a novel ubiquitin ligase (E3) involved in the regulation of carbohydrate metabolism. Mol Biol Cell 19: 3323–3333.

35. Qiao S, Langlois CR, Chrustowicz J, Sherpa D, Karayel O, Hansen FM, Beier V, Gronau von S, Bollschweiler D, Schäfer T, et al. (2020) Interconversion between Anticipatory and Active GID E3 Ubiquitin Ligase Conformations via Metabolically Driven Substrate Receptor Assembly. Mol Cell 77: 150–163.e159.

36. Sherpa D, Chrustowicz J, Qiao S, Langlois CR, Hehl LA, Gottemukkala KV, Hansen FM, Karayel O, Gronau von S, Prabu JR, et al. (2021) GID E3 ligase supramolecular chelate assembly configures multipronged ubiquitin targeting of an oligomeric metabolic enzyme. Mol Cell 81: 2445–2459.e13.

37. Menssen R, Schweiggert J, Schreiner J, Kušević D, Reuther J, Braun B, Wolf DH (2012) Exploring the Topology of the Gid Complex, the E3 Ubiquitin Ligase Involved in Catabolite-induced Degradation of Gluconeogenic Enzymes. J Biol Chem 287: 25602–25614.

38. Kong K-YE, Fischer B, Meurer M, Kats I, Li Z, Rühle F, Barry JD, Kirrmaier D, Chevyreva V, San Luis B-J, et al. (2021) Timer-based proteomic profiling of the ubiquitin-proteasome system reveals a substrate receptor of the GID ubiquitin ligase. Mol Cell 81: 2460–2476.e11.

39. Melnykov A, Chen S-J, Varshavsky A (2019) Gid10 as an alternative N-recognin of the Pro/N-degron pathway. Proc Natl Acad Sci USA 116: 15914–15923.

40. Karayel O, Michaelis AC, Mann M, Schulman BA, Langlois CR (2020) DIA-based systems biology approach unveils E3 ubiquitin ligase-dependent responses to a metabolic shift. Proc Natl Acad Sci USA 117: 32806–32815.

41. Oh J-H, Chen S-J, Varshavsky A (2017) A reference-based protein degradation assay without global translation inhibitors. J Biol Chem 292: 21457–21465.

42. Gasch AP, Spellman PT, Kao CM, Carmel-Harel O, Eisen MB, Storz G, Botstein D, Brown PO (2000) Genomic expression programs in the response of yeast cells to environmental changes. Mol Biol Cell 11: 4241–4257.

43. Menssen R, Bui K, Wolf DH (2018) Regulation of the Gid ubiquitin ligase recognition subunit Gid4. FEBS Lett 592: 3286–3294.

44. Lu J, Kobayashi R, Brill SJ (1996) Characterization of a high mobility group 1/2 homolog in yeast. J Biol Chem 271: 33678–33685.

45. Lussier M, White AM, Sheraton J, di Paolo T, Treadwell J, Southard SB, Horenstein CI, Chen-Weiner J, Ram AF, Kapteyn JC, et al. (1997) Large scale identification of genes involved in cell surface biosynthesis and architecture in Saccharomyces cerevisiae. Genetics 147: 435–450.

46. Dong C, Zhang H, Li L, Tempel W, Loppnau P, Min J (2018) Molecular basis of GID4-mediated recognition of degrons for the Pro/N-end rule pathway. Nat Chem Biol 14: 466–473.

47. Dong C, Chen S-J, Melnykov A, Weirich S, Sun K, Jeltsch A, Varshavsky A, Min J (2020) Recognition of nonproline N-terminal residues by the Pro/N-degron pathway. Proc Natl Acad Sci USA 117: 14158–14167.

48. Swaney DL, Beltrao P, Starita L, Guo A, Rush J, Fields S, Krogan NJ, Villén J (2013) Global analysis of phosphorylation and ubiquitylation cross-talk in protein degradation. Nat Methods 10: 676–682.

49. Babst M (2020) Regulation of nutrient transporters by metabolic and environmental stresses. Curr Opin Cell Biol 65: 35–41.

50. Negoro H, Matsumura K, Matsuda F, Shimizu H, Hata Y, Ishida H (2020) Effects of mutations of GID protein-coding genes on malate production and enzyme expression profiles in Saccharomyces cerevisiae. Appl Microbiol Biotechnol 104: 4971–4983.

51. Baile MG, Guiney EL, Sanford EJ, MacGurn JA, Smolka MB, Emr SD (2019) Activity of a ubiquitin ligase adaptor is regulated by disordered insertions in its arrestin domain. Mol Biol Cell 30: 3057–3072.

52. MacDonald C, Winistorfer S, Pope RM, Wright ME, Piper RC (2017) Enzyme reversal to explore the function of yeast E3 ubiquitin-ligases. Traffic 18: 465–484.

53. Maspero E, Mari S, Valentini E, Musacchio A, Fish A, Pasqualato S, Polo S (2011) Structure of the HECT:ubiquitin complex and its role in ubiquitin chain elongation. EMBO Rep 12: 342–349.

54. Kim HC, Steffen AM, Oldham ML, Chen J, Huibregtse JM (2011) Structure and function of a HECT domain ubiquitin-binding site. EMBO Rep 12: 334–341.

55. Brown CR, Dunton D, Chiang H-L (2010) The vacuole import and degradation pathway utilizes early steps of endocytosis and actin polymerization to deliver cargo proteins to the vacuole for degradation. J Biol Chem 285: 1516–1528.

56. Giardina BJ, Dunton D, Chiang H-L (2013) Vid28 protein is required for the association of vacuole import and degradation (Vid) vesicles with actin patches and the retention of Vid vesicle proteins in the intracellular fraction. J Biol Chem 288: 11636–11648.

57. Horak J, Regelmann J, Wolf DH (2002) Two Distinct Proteolytic Systems Responsible for Glucose-induced Degradation of Fructose-1,6-bisphosphatase and the Gal2p Transporter in the Yeast Saccharomyces cerevisiaeShare the Same Protein Components of the Glucose Signaling Pathway. J Biol Chem 277: 8248–8254.

58. Herst PM, Perrone GG, Dawes IW, Bircham PW, Berridge MV (2008) Plasma membrane electron transport in Saccharomyces cerevisiae depends on the presence of mitochondrial respiratory subunits. FEMS Yeast Research 8: 897–905.

59. Salemi LM, Maitland MER, McTavish CJ, Schild-Poulter C (2017) Cell signalling pathway regulation by RanBPM: molecular insights and disease implications. Open Biol 7:.

60. Lampert F, Stafa D, Goga A, Soste MV, Gilberto S, Olieric N, Picotti P, Stoffel M, Peter M (2018) The multi-subunit GID/CTLH E3 ubiquitin ligase promotes cell proliferation and targets the transcription factor Hbp1 for degradation. Elife 7:.

61. Liu H, Ding J, Köhnlein K, Urban N, Ori A, Villavicencio-Lorini P, Walentek P, Klotz L-O, Hollemann T, Pfirrmann T (2020) The GID ubiquitin ligase complex is a regulator of AMPK activity and organismal lifespan. Autophagy 16: 1618–1634.

62. Knop M, Siegers K, Pereira G, Zachariae W, Winsor B, Nasmyth K, Schiebel E (1999) Epitope tagging of yeast genes using a PCR-based strategy: more tags and improved practical routines. Yeast 15: 963–972.

63. Janke C, Magiera MM, Rathfelder N, Taxis C, Reber S, Maekawa H, Moreno-Borchart A, Doenges G, Schwob E, Schiebel E, et al. (2004) A versatile toolbox for PCR-based tagging of yeast genes: new fluorescent proteins, more markers and promoter substitution cassettes. Yeast 21: 947–962.

64. Storici F, Resnick MA (2003) Delitto perfetto targeted mutagenesis in yeast with oligonucleotides. Genet Eng (N Y) 25: 189–207.

65. Gibson DG, Young L, Chuang R-Y, Venter JC, Hutchison CA, Smith HO (2009) Enzymatic assembly of DNA molecules up to several hundred kilobases. Nat Methods 6: 343–345.

66. Weissmann F, Petzold G, VanderLinden R, Huis In ‘t Veld PJ, Brown NG, Lampert F, Westermann S, Stark H, Schulman BA, Peters J-M (2016) biGBac enables rapid gene assembly for the expression of large multisubunit protein complexes. Proc Natl Acad Sci USA 113: E2564–E2569.

67. Adams PD, Afonine PV, Bunkóczi G, Chen VB, Davis IW, Echols N, Headd JJ, Hung L-W, Kapral GJ, Grosse-Kunstleve RW, et al. (2010) PHENIX: a comprehensive Python-based system for macromolecular structure solution. Acta Crystallogr D Biol Crystallogr 66: 213–221.

68. Afonine PV, Poon BK, Read RJ, Sobolev OV, Terwilliger TC, Urzhumtsev A, Adams PD (2018) Real-space refinement in PHENIX for cryo-EM and crystallography. Acta Crystallogr D Struct Biol 74: 531–544.

69. DiMaio F, Echols N, Headd JJ, Terwilliger TC, Adams PD, Baker D (2013) Improved low-resolution crystallographic refinement with Phenix and Rosetta. Nat Methods 10: 1102–1104.

70. Emsley P, Cowtan K (2004) Coot: model-building tools for molecular graphics. Acta Crystallogr D Biol Crystallogr 60: 2126–2132.

71. Emsley P, Lohkamp B, Scott WG, Cowtan K (2010) Features and development of Coot. Acta Crystallogr D Biol Crystallogr 66: 486–501.

72. Tyanova S, Temu T, Sinitcyn P, Carlson A, Hein MY, Geiger T, Mann M, Cox J (2016) The Perseus computational platform for comprehensive analysis of (prote)omics data. Nat Methods 13: 731–740.

